# OMA1 protease eliminates arrested protein import intermediates upon depolarization of the inner mitochondrial membrane

**DOI:** 10.1101/2023.06.08.543713

**Authors:** Magda Krakowczyk, Anna M. Lenkiewicz, Tomasz Sitarz, Ben Hur Marins Mussulini, Vanessa Linke, Dominika Malinska, Andrzej Szczepankiewicz, Agata Wydrych, Hanna Nieznanska, Remigiusz A. Serwa, Agnieszka Chacinska, Piotr Bragoszewski

**Affiliations:** Nencki Institute of Experimental Biology, Polish Academy of Sciences, Warsaw, Poland; Centre of New Technologies, University of Warsaw, Warsaw, Poland; IMol Polish Academy of Sciences, Warsaw, Poland; ReMedy International Research Agenda Unit, IMol Polish Academy of Sciences, Warsaw, Poland

## Abstract

Most mitochondrial proteins originate from the cytosol and require active transport into the organelle. Such precursor proteins must be largely unfolded to pass through translocation channels in mitochondrial membranes. Misfolding of transported proteins can result in their arrest and translocation failure. Arrested proteins block further import, disturbing mitochondrial functions and cellular proteostasis. Cellular responses to translocation failure have been defined in yeast. To discover molecular mechanisms that resolve failed import events in human cells, we developed the translocase clogging model using a fusion protein with a rigid domain. The mechanism we uncover differs significantly from these described in fungi, where ATPase-driven extraction of blocked protein is directly coupled with proteasomal processing. We found human cells to rely primarily on mitochondrial factors to clear translocation channel blockage. The mitochondrial membrane depolarization triggered proteolytic cleavage of the stalled protein, which involved mitochondrial protease OMA1. The cleavage allowed releasing the protein fragment that blocked the translocase. The released fragment was further cleared in the cytosol by the valosin containing protein (VCP)/p97 and proteasome.

## Introduction

Cellular proteomes are composed of thousands of different proteins, ensuring the diversity of the molecular functions and the cell’s metabolic flexibility. Biogenesis and maintenance of diverse proteomes are challenging tasks. Cellular proteins must be produced with a correct amino acid sequence, transported to their specific destination, and folded into mature, functional structures. Failure of any of these steps not only results in the formation of erroneous proteins but has the capacity to disturb cellular protein homeostasis. This necessitates quality control (QC) mechanisms that repair or remove aberrant proteins and restore homeostasis.

Protein transport is a critical aspect in the biogenesis of cellular organelles with sharply defined boundaries. This is apparent for mitochondria, with two specialized membrane systems, the outer mitochondrial membrane (OM) and the inner mitochondrial membrane (IM), sculpting their architecture. These membranes envelop aqueous sub-compartments called the mitochondrial matrix and the intermembrane space (IMS). The architecture of mitochondria allows these organelles to embody a unique environment and harness vital metabolic processes, including oxidative phosphorylation. The proteome of human mitochondria comprises well over a thousand diverse proteins (Rath et al., 2021; Morgenstern et al., 2021). Nearly all of these are produced in the cytosol as protein precursors, which require selective and accurate transport to their destination sites within the organelle. Targeting information that is embedded in their amino acid sequence induces their interaction with the mitochondrial protein import machinery (Becker et al., 2019; Wasilewski et al., 2016; Bykov et al., 2020).

The diversity of mitochondrial proteins and their sub-mitochondrial locations are reflected by the diversity of their mitochondrial targeting signals (MTS) and sorting mechanisms. Still, the primary step of the import is common to most precursors. They cross the OM via the translocase of the outer membrane (TOM) complex that serves as a passage, precursor receptor, and import regulator (Wasilewski et al., 2016; Becker et al., 2019). Protein-conducting channels across the OM are formed by the β-barrel protein Tom40, a central component of the TOM complex. Tom40 proteins are linked and organized by Tom22 molecules and small TOM subunits: Tom5, Tom6, and Tom7. Tom20 and Tom70 subunits expose receptor domains to recognize precursors and cooperate with Tom22 to initiate the import.

After crossing the OM via TOM, proteins are routed to their final destinations by several protein-sorting and -assembly pathways. The largest group of precursors follows the presequence pathway operated by the translocase of the inner membrane, the TIM23 complex. Such precursors possess N-terminal cleavable MTS sequences and rely on the IM proton gradient to initiate and assist their translocation, which is later taken over by the presequence translocase-associated motor (PAM) (Schendzielorz et al., 2017). TIM23 substrates are fully translocated into the matrix or laterally released for IM integration. Upon maturation, substrates of the presequence pathway undergo proteolytic removal of the MTS (Mossmann et al., 2012; Vogtle et al., 2009). Other protein translocation pathways include TIM22 for IM integration of multiple membrane-spanning substrates, the sorting and assembly machinery (SAM), or the insertase of the mitochondrial outer membrane (MIM) for integration into the OM, and the mitochondrial import and assembly (MIA) pathway which operates in the IMS (Wasilewski et al., 2016; Becker et al., 2019).

Protein import routes within mitochondria differ by their driving force and throughput (Schäfer et al., 2022; Morgenstern et al., 2017). Import efficacy also depends on individual precursors’ characteristics. Common is that precursor folding occurs after they reach their destination. Proteins directed to mitochondria must be unfolded while passing the translocases since the lumen of the protein-conducting channel formed by Tom40 protein would not accommodate most folded domains (Wiedemann and Pfanner, 2017; Kater et al., 2020; Tucker and Park, 2019; Ahting et al., 2001). Thus, a folded domain present in a precursor can stall during import, preventing the completion of the translocation (Schleyer and Neupert, 1985; Rassow et al., 1989; Wienhues et al., 1991; Glick et al., 1993; Kanamori et al., 1997; Schulke et al., 1997; Gaume et al., 1998; Schulke et al., 1999; Voisine et al., 1999; Chacinska et al., 2003; Gold et al., 2014). The TOM channel allows the backward movement of proteins, so arrested precursors can be released back into the cytosol (Bragoszewski et al., 2015). However, precursors that stall during import can span both outer and inner membranes in a stable complex with TOM and TIM translocases (Chacinska et al., 2003; Gomkale et al., 2021). Such stabilization prevents further mobility of the arrested protein. Substrates of both TIM23 and TIM22 could be affected by such arrests (Shiota et al., 2015). *In vitro* studies with synthetic precursor proteins provided the first evidence that arrested import intermediates sequester available translocases. Studies in yeast revealed that translocase blocking results in a growth defect (Wienhues et al., 1991; Schulke et al., 1997, 1999). The growth impairment reflects the severity of the issue, as the impact of protein import blockage goes beyond disturbing mitochondria’s bioenergetic and metabolic functions. Unimported mitochondrial proteins accumulate in the cytosol, increasing the risk of misfolding and aggregation (Nowicka et al., 2021). Importantly, native precursor proteins were found to stall during import (Weidberg and Amon, 2018; Boos et al., 2019; Glick et al., 1993).

Given the potentially severe consequences of clogging, efficient response mechanisms are necessary. As a first tier of defense against precursor proteins mislocalization, yeast cells were found to increase proteasome assembly and activity (Wrobel et al., 2015). Proteasomal capacity is further boosted by upregulated expression of proteasome components, a part of a broader transcriptional program named mitoprotein-induced stress response that also upregulates chaperones (Boos et al., 2019). Simultaneously, transcripts encoding multiple mitochondrial proteins are downregulated, decreasing the load on mitochondrial translocases (Boos et al., 2019). In parallel with increased degradation, general protein synthesis becomes attenuated to stop the further build-up of mislocalized proteins (Wang and Chen, 2015; Wrobel et al., 2015). Thus, the failure of mitochondrial import can globally alter both sides of the protein turnover cycle, namely protein synthesis and degradation.

Resolving a translocase blockage requires the removal of the stalled cargo. Machinery that extracts proteins and targets them for degradation was first discovered at endoplasmic reticulum membranes and termed ER-associated protein degradation (ERAD) (Romisch, 2005; Tsai et al., 2002; Hirsch et al., 2009), followed by observation of a similar mechanism operating on mitochondria termed mitochondria-associated degradation (MAD) (Tanaka et al., 2010; Heo et al., 2010; Xu et al., 2011). A closely related pathway, termed mitochondrial protein translocation-associated degradation (mitoTAD), was shown to clear precursors that stall in mitochondrial translocons of *S. cerevisiae* (Martensson et al., 2019). The above processes share a dependence on a ubiquitin-dependent chaperone complex built around valosin containing protein AAA ATPase (VCP, also known as p97 or Cdc48 in yeast). The ATPase activity of Cdc48 allows extraction of the proteins which are subsequently degraded by the proteasome. Surveillance by mitoTAD is constitutively active, assuring import capacity in normal conditions. An additional, inducible mechanism that specifically reacts to translocase overload by the IM-targeted proteins was also discovered in yeast and termed mitochondrial compromised protein import response (mitoCPR) (Weidberg and Amon, 2018). In this inducible mechanism, another AAA ATPase – Msp1 is recruited to extract the arrested precursor, allowing its further degradation by the proteasome (Weidberg and Amon, 2018; Basch et al., 2020).

Several molecular mechanisms responding to mitochondrial protein translocase clogging reported to date show that cells are equipped with redundant defense paths, highlighting the biological significance of the problem. However, our understanding of these QC mechanisms originates vastly from studies that used yeast as a research model. To understand how the mechanisms discovered in fungi translate to higher eukaryotes, we investigate translocase clogging in human cells. In this work, we developed and validated a cellular model of translocase clogging based on stably folding GFP protein (i.e., superfolder GFP; sfGFP). The sfGFP tag within the mitochondria-targeted fusion caused its arrest in the TOM channel, interfering with the import of other proteins and disturbing mitochondrial functions. Using this model, we report a significant role of mitochondrial proteases in clearing arrested precursors that block mitochondrial translocases. Surprisingly, despite its negative impact, the arrested protein proved stable in a steady state and was rapidly cleared only upon dissipation of the inner membrane potential. The depolarization triggered proteolytic cleavage of the arrested precursor allowing the release of the unimported fragment from the translocase and restoration of mitochondrial architecture. The released protein became a substrate of cytosolic degradation, mediated by VCP and proteasome. We conclude that in human mitochondria, precursors in transit may become stably bound and require intramitochondrial cleavage for effective release. The observed mechanism differs from those described in fungi, where AAA ATPases are recruited to extract arrested cargo. Also, the role of the proteasome is less direct as the initial cleavage of arrested protein is executed by mitochondrial proteases, including OMA1.

## Results

### Model in human cells

To study the quality control (QC) of mitochondrial protein translocation, we established an inducible import failure model in a human cell line. We based the model on our earlier finding that tagging mitochondrial proteins with the tandem fluorescent protein timer (tFT; a fusion of two fluorescent proteins: mCherry and superfolder GFP (sfGFP)) interferes with their import (Kowalski et al., 2018). Import disturbance likely could be attributed to sfGFP’s resistance to unfolding (Khmelinskii et al., 2016). This agrees with a recent study showing that sfGFP stalls in the mitochondrial translocation machinery of yeast if fused to the N-terminal targeting sequence (Gomkale et al., 2021). Here we designed a precursor protein consisting of the ATP5MG fused to a C-terminal ‘clogging’ tFT tag (Fig. 1A) for expression in human cells. We used the Flp-In T-REx HEK-293 cell line, which allowed for recombination-based stable integration of the transgene and for its inducible expression upon tetracycline addition to the culture media (Fig. 1B). Cells expressing ATP5MG-tFT fusion accumulated a full-size fusion protein but also its smaller fragments which were detected by an anti-GFP antibody (Fig. 1B, C).

**Fig. 1.**
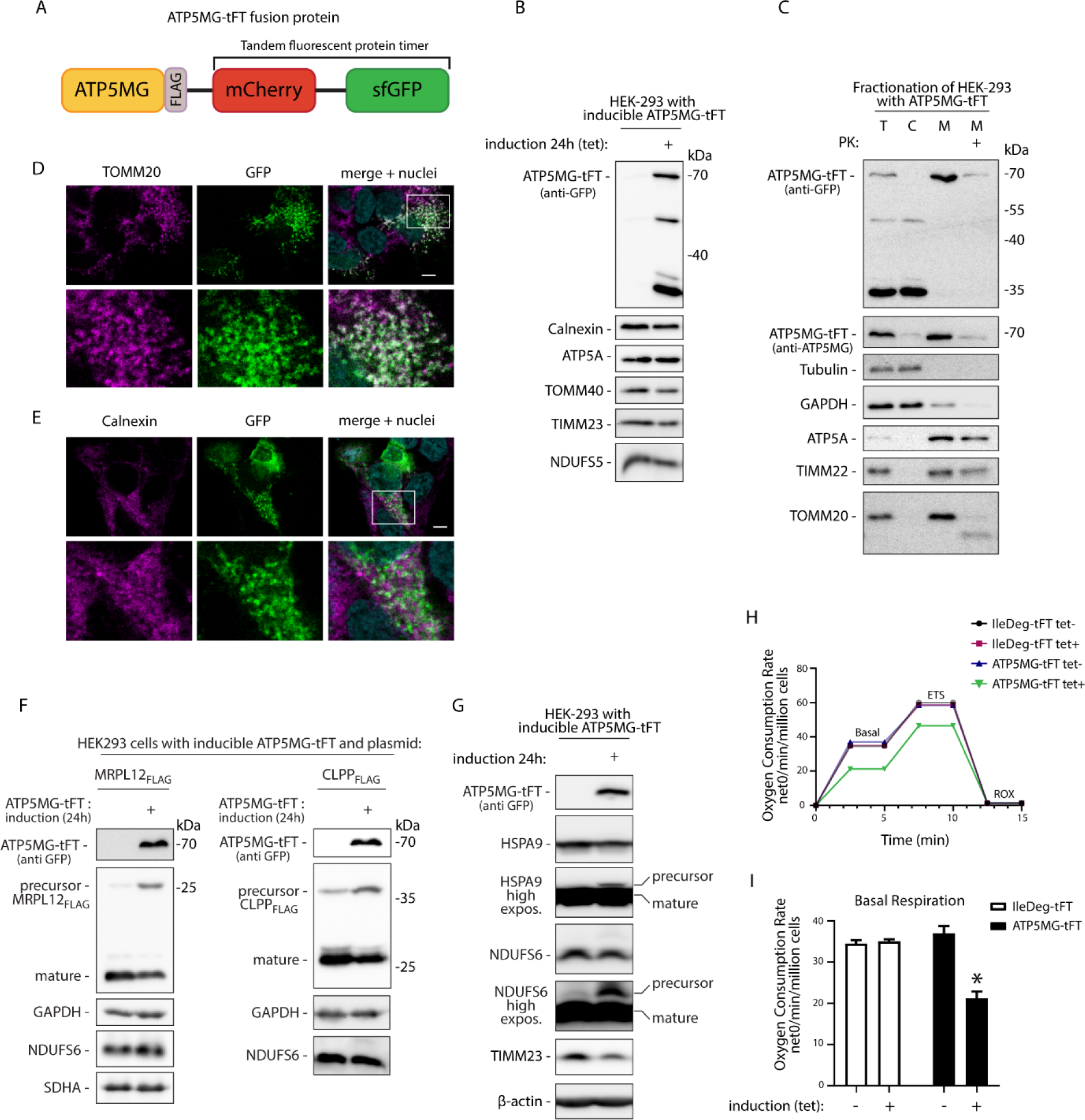
ATP5MG-tFT fusion causes mitochondrial protein translocase clogging and impairs mitochondrial protein import. (A) Schematic representation of ATP5MG-tFT fusion protein used to model translocase blocking. ATP5MG was expressed in fusion with the tandem fluorescent protein timer (tFT) consisting of mCherry and super folder GFP (sfGFP) proteins. (B) Tetracycline-inducible expression of ATP5MG-tFT in HEK-293 cells. The cells were grown with or without the addition of tetracycline (1 µg/mL) for 24 h. Total protein extracts were analyzed with indicated antibodies. Expression of the fusion protein was tested with anti-GFP antibody. (C) Subcellular fractionation of HEK-293 cells expressing ATP5MG-tFT protein. After differential centrifugation, samples were analyzed by SDS–PAGE and western blotting. T-total, C-cytoplasm, M-mitochondria, PK-proteinase K treatment (25 µg/mL). (D, E) Intracellular distribution of ATP5MG-tFT in HEK-293 cells visualized by confocal imaging. ATP5MG-tFT was detected by GFP fluorescence. Cells were immunolabeled using antibodies against TOMM20 (D) or calnexin (E), to visualize mitochondria and ER, respectively. Cell nuclei were stained with DAPI. Scale bars 5 μm. Bottom panels show enlarged fragment marked in the top panel. (F) Co-expression of ATP5MG-tFT and plasmid-encoded CLPP_FLAG_ or MRPL12_FLAG_ proteins. Total cell extracts were analyzed by SDS–PAGE and western blotting. Precursor and mature forms processed in mitochondria were detected by anti-FLAG antibodies. (G) Cells with or without induction of ATP5MG-tFT were analyzed as in (B). High exposure images for HSPA9 and NDUFS6 western blots are shown. (H) Oxygen consumption rate (OCR) in cells expressing different tFT fusions. ETS (maximal respiration), ROX (residual oxygen consumption). (I) Basal respiration quantified from H. OCR was normalized to the number of cells. Data are presented as the mean ± SEM (n = 3); *p<0.05 (two-way ANOVA followed by multicomparision test).

Efficient targeting into mitochondria is a prerequisite for a fusion protein to stall in the translocation machinery. Accordingly, ATP5MG-tFT protein was recovered in the mitochondrial fraction, while faster-migrating fragments were located in the cytosol (Fig. 1C). The clogging fusion was susceptible to externally added protease, as was the outer membrane (OM) protein TOMM20 but not the internal mitochondrial proteins (ATP5A, TIMM22) (Fig. 1C). This indicates that at least part of the fusion was exposed on the outer side of the OM.

We confirmed the sub-cellular localization of the ATP5MG-tFT by confocal microscopy. The sfGFP fluorescent signal colocalized with the OM marker protein TOMM20 but not with the endoplasmic reticulum (ER) marker calnexin (Fig. 1D, E). The sfGFP signal also colocalized with the mitochondrial stain Mitotracker Deep Red (Fig. S1A). In this case, GFP fluorescence partially surrounded a more centered Mitotracker signal, which matches their expected localization at the OM and inside the matrix.

To verify that the fusion protein blocks mitochondrial translocases, we tested its effect on the import of other mitochondrial proteins. We co-expressed ATP5MG-tFT together with mitochondria-targeted proteins CLPP_FLAG_ or MRPL_FLAG_. Both of these proteins contain relatively large N-terminal MTS signals, which make up ∼20% of their molecular mass and are cleaved off upon maturation in the matrix (Calvo et al., 2017). Thus, mature and precursor forms can be easily separated by electrophoresis. Indeed, upon the expression of CLPP_FLAG_ and MRPL_FLAG_ in HEK293 cells, well-separated bands could be detected with anti-FLAG antibodies, corresponding to precursor and mature forms (Fig. 1F). In standard conditions, CLPP_FLAG_ or MRPL_FLAG_ were effectively imported and processed in mitochondria, as manifested by the accumulation of smaller mature forms of these proteins. However, when expressed along ATP5MG-tFT fusion, we observed decreased accumulation of mature forms and a noticeable increase in unprocessed precursors (Fig. 1F).

We also looked for the accumulation of unprocessed precursor forms of native mitochondrial proteins. In physiological conditions, cellular levels of most mitochondrial protein precursors are low and often undetectable by western blot. However, translocase blocking can lead to the accumulation of unimported proteins (Weidberg and Amon, 2018; Boos et al., 2019; Martensson et al., 2019). Indeed, longer exposure of western blot images revealed the accumulation of higher molecular mass forms of mitochondrial heat shock protein HSPA9 and NDUFS6 (Fig. 1G). Such slower migrating forms, likely unprocessed precursors, were detected only in the cells expressing ATP5MG-tFT but not in control cells (i.e., without induction of expression). Together, our observations provided evidence for a mitochondrial protein import defect related to the expression of the model protein.

To further characterize the model cells, we evaluated oxygen consumption rates (OCRs) (Fig. 1H). Translocase blocking in yeast was demonstrated to cause a general decline in mitochondrial respiratory function. To check whether our fusion protein induces a similar defect, we compared oxygen utilization in untreated cells (i.e., without expression induction of ATP5MG-tFT) or 24 hours after inducing ATP5MG-tFT production. We also included a cell line that expresses tFT fusion not directed to mitochondria (IleDeg-tFT) as a control. The expression of IleDeg-tFT did not affect any of the measured respiratory parameters. However, cells expressing the mitochondria-directed fusion displayed significantly lower basal respiration (Fig. 1I) and a tendency towards decreased maximum respiration (Fig. S1C).

The expression of ATP5MG-tFT clogging fusion decreased cell proliferation. The number of cells in the population after 48 hours of expression induction was 79% of the population without induction. The decrease was specific to the clogging fusion as it was not observed in control cells. Expression of IleDeg-tFT fusion did not affect cell numbers. Similarly, HEK-293 Flp-In cells with no inducible transgene remained unaffected by inducing agent treatment (1 µg/ml tetracycline). The decrease in ATP5MG-tFT cell numbers was significant compared to both, IleDeg-tFT expressing and empty HEK-293 Flp-In cells (Fig. S1D). Still, the proliferation reduction appears mild compared to severe growth defects reported in yeast (Boos et al., 2019). This possibly reflects modest fusion protein expression levels characteristic of Flp-In T-REx cells.

Together, we found that the model fusion was directed to mitochondria, which interfered with the import of other proteins, disturbed mitochondrial functions, and impacted cell proliferation. These properties substantiate the probability of translocase clogging by ATP5MG-tFT protein.

### Molecular interactions of the clogging protein validate translocase blocking

To define binding partners of the model fusion, we have developed a co-immunoprecipitation (Co-IP) protocol. We used GFP-Trap beads to purify ATP5MG-tFT from total cell extracts (Fig. 2A). The depletion of the bait in unbound fractions confirmed the high IP efficiency. Using antibodies against TOMM40 and TOMM20 proteins, we confirmed that the model protein interacted with the TOM translocase. This membrane complex was co-purified only with ATP5MG-tFT, but not with the control tFT fusion (IleDeg-tFT) or from the cells with no GFP. Interestingly, we did not observe similar copurification of TIMM23 protein of the inner membrane translocase. Instead, we found the ATP5A protein of the ATP synthase complex among proteins co-purified with the clogging fusion. This indicates that the N-terminal part of the fusion likely reached its destination.

**Fig. 2.**
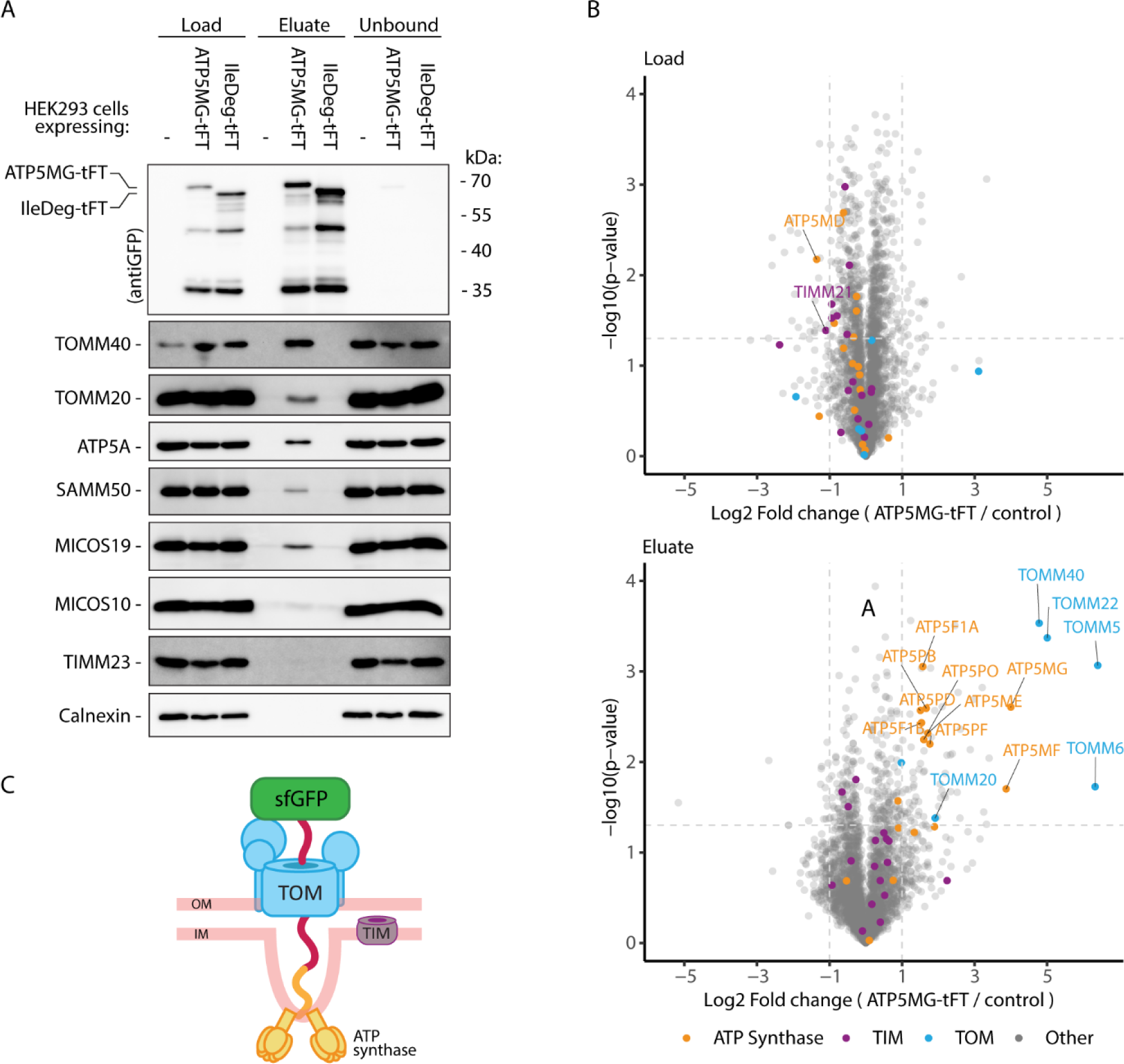
Mitochondria-directed tFT fusion protein interacts with mitochondrial protein translocation channel. (A) Lysates of HEK293 cells expressing ATP5MG-tFT, or ILE-Deg-tFT (i.e., tFT fusion not directed to mitochondria), and control cells with no transgene were immunoprecipitated with anti-GFP beads. Proteins from the initial sample (Load, 10%), the Eluate (100%), and the sample remaining after IP (Unbound, 10%) were analyzed using SDS–PAGE and immunoblotting with the indicated antibodies. (B) Proteins from cellular lysates (Load) and purified with anti-GFP beads (Eluate) were analyzed by LC-MS/MS (n=4). In volcano plots, the x-axis represents the log_2_ fold change of protein levels in ATP5MG-tFT samples compared to control cells samples. Total lysates (top) and eluates (bottom) were analyzed. Orange, magenta, and blue indicate proteins belonging to ATP synthase, TIM, and TOM complexes, respectively; of these significantly changed proteins (Student’s t-test p<0.05, |log_2_ fold change| >1) were labeled. (C) Schematic depiction of ATP5MG-tFT plausible topology, interacting with ATP synthase and TOM translocase.

Having established an effective Co-IP protocol, we aimed to define the interactome of the clogging protein using mass spectrometry (MS)-based proteomics. The experiments compared cells expressing the clogging protein and control cells not expressing any fusion protein (Fig. 2B). Additionally, we compared the clogging model to non-mitochondrial tFT fusion IleDeg-tFT (Fig. S2A, B). We analyzed co-IP eluates and corresponding total protein extracts. Out of 3063 identified proteins, 86 were significantly enriched in clogger IP samples. Among the most enriched were components of the mitochondrial import machinery, namely TOM translocase subunits (Fig. 2B bottom and Fig. S2B). Although ATP5MG directs the clogging construct for IM integration, we did not detect enrichment of the components of the IM translocases TIM23 or TIM22. Instead, subunits of the ATP synthase were among the significantly enriched genes. This indicates that the N-terminal part of ATP5MG-tFT could complete its import, becoming integrated into the ATP synthase complex. Together with the robust interaction with TOM components, these results signify stable OM translocase clogging by the fusion. Based on a recently published cryo-EM structure of the mammalian ATP synthase, we expect ATP5MG to be located mainly within the membrane and with its C-terminus exposed on the IMS side of the IM (Spikes et al., 2020). Combining this data, we propose the topology of the arrested protein as outlined in Fig. 2C.

Other identified interactors of the fusion protein could not be grouped by molecular function (see Supplementary Table 1 for the complete list). Next, a comparative analysis of protein changes in cell lysates that were inputs for the co-IP experiment was performed (Fig. 2B, see Load). This served to ensure that the expression of fusion proteins alone did not cause changes in the expression of other proteins. Indeed, only a few proteins showed a differential expression in lysates originating from ATP5MG-tFT-expressing vs. control cells, none of which was identified as an ATP5MG-tFT binder. Importantly, none of the ATP synthase or TOM components were upregulated in ATP5MG-tFT cells supporting the specificity of the IP enrichment. We conclude that short-time expression of the clogging fusion (24 h of induction prior to MS analysis) does not trigger substantial proteome remodeling.

### The stability of the clogging fusion depends on the mitochondrial membrane potential

Blocking of the translocase by a stalled cargo disturbs mitochondrial biogenesis but also cytosolic proteostasis due to the increasing load of mislocalized proteins. These substantiate the need for effective QC. Studies in yeast found proteins that stall in mitochondrial translocases to be destabilized by the action of proteolytic QC machinery. We tested the stability of ATP5MG-tFT fusion using cycloheximide to block cellular protein synthesis (Fig. 3A). Surprisingly, we did not observe substantial degradation of the clogging fusion protein over eight hours. This indicated that despite being blocked in the TOM channel, ATP5MG-tFT remained stable in basal conditions. However, once we dissipated mitochondrial membrane potential using CCCP, the full-size fusion protein was effectively removed from the cells (Fig. 3B). The rapid removal of the clogging protein was specific to the membrane potential dissipation. Accordingly, treatment with BAM15 protonophore, which unlike CCCP is more selective to the mitochondrial membrane as it does not depolarise the cellular membrane (Kenwood et al., 2014), resulted in degradation of the arrested fusion protein (Fig. 3C). Another ionophore, valinomycin, which also efficiently depolarizes mitochondria, impacted the accumulation of the ATP5MG-tFT similarly to CCCP and BAM15 (Fig. 3D). At the same time, mitochondrial poisons that have weaker or no effect on mitochondrial membrane potential: oligomycin A, antimycin A, and rotenone, did not cause clogging fusion destabilization.

**Fig. 3.**
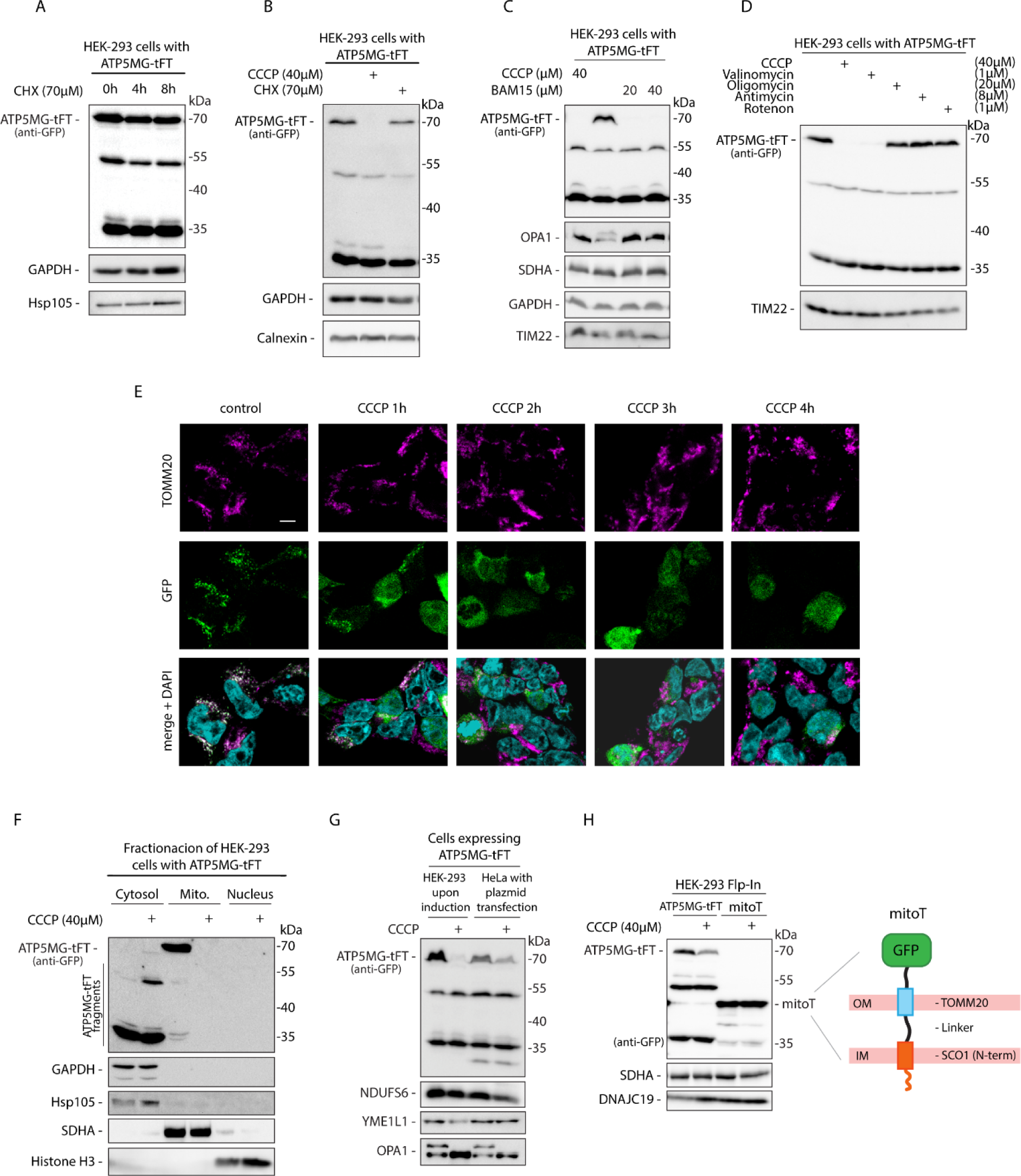
Protein blocked in the translocase becomes destabilized upon dissipation of Mitochondrial membrane potential. (A, B, C, D) Cellular levels of ATP5MG-tFT protein in HEK-293 cells following indicated treatments: (A) cells were treated with cycloheximide (CHX, 70 µM) to inhibit protein synthesis for the specified time; (B) cells were treated with CCCP(40 µM) to dissipate mitochondrial membrane potential; (C) cells were treated with indicated concentrations of protonophores CCCP or BAM15 for 2 h; (D) cells were treated for 2 h with CCCP, valinomycin, oligomycin, antimycin A, or rotenone at the indicated concentrations; (E) Confocal imaging of HEK293 cells expressing ATP5MG-tFT (GFP signal) treated with 40 µM CCCP for the indicated time. Mitochondria were visualized by TOMM20 immunostaining; cell nuclei were stained with DAPI. Scale bar: 5 μm. (F) Impact of 2 h of CCCP treatment on the presence of ATP5MG-tFT in cytosolic, mitochondrial (Mito.), and nuclear fractions of HEK-293 cells. Fractions were analyzed by western blotting using anti-GFP antibody to detect ATP5MG-tFT. For validation, cytosolic proteins GAPDH and Hsp105, mitochondrial marker SDHA, and nuclear marker Histone H3, were detected. (G) HEK-293 (after tetracycline induction) and HeLa (plasmid transfected) cells expressing ATP5MG-tFT were treated or not treated with 40 µM CCCP for the final 2 h of the culture. Total protein extracts were analyzed by SDS-PAGE and western blotting. Expression of the fusion proteins was tested with an anti-GFP antibody. (H) HEK-293 cells were transfected with mitoT (Viana et al., 2021) or ATP5MG-tFT expression vectors. CCCP treatment was applied for 2 hours. Total protein extracts were analyzed by SDS-PAGE and western blotting. Expression of the fusion proteins was tested with an anti-GFP antibody; Schematic depiction of the mitoT membrane tether construct (Viana et al., 2021).

We have monitored the uncoupling-induced changes of the GFP signal (i.e., part of the tFT fusion) by confocal imaging at different times after CCCP addition (Fig. 3E). Before treatment, GFP fluorescence was distributed in defined foci that colocalized with the mitochondrial marker TOMM20. After one hour of CCCP treatment, the fluorescent signal became diffused, while after two hours, GFP appeared evenly distributed in the cytosol, showing no apparent colocalization with TOMM20. During more prolonged CCCP treatment, the GFP signal gradually diminished, with the remaining fluorescence detectable mainly in the cells’ nuclei.

To gain an additional perspective on how depolarization-induced degradation affects the localization of fusion protein and its fragments, we analyzed their presence in cytosolic, mitochondrial, and nuclear fractions (Fig. 3F). We detected full-size clogging fusion only in the mitochondria-enriched fraction. Upon 2 hours of treatment with CCCP, this band was depleted from mitochondria and was not recovered in any other fraction. However, CCCP treatment increased the level of lower molecular mass fusion fragments in the cytosol, which we consider degradation products. No GFP-related signal was detected in the nuclear fraction.

To verify that depolarization-activated degradation is not specific to HEK-293 cells, we transiently expressed ATP5MG-tFT clogging fusion in HeLa cells. Both cell lines responded to uncoupler treatment as indicated by OPA1 processing (Baker et al., 2014; Zhang et al., 2014). Similarly, we observed a full-size protein ATP5MG-tFT level reduction in both cell types when adding CCCP (Fig. 3G).

Being threaded through the TOM complex and bound with ATP synthase, ATP5MG-tFT connects two mitochondrial membranes. We asked if protein connecting two membranes but not stuck in the translocase would also be processed upon depolarization. To test this, we expressed mitoT tether fusion, which was designed to span two membranes (Viana et al., 2021). MitoT also includes GFP moiety, which is preceded by a transmembrane fragment of TOMM20 protein. Thus the GFP of mitoT remains outside the organelle, not colliding with the fusion’s import (Fig. 3H, see scheme). In contrast to ATP5MG-tFT, mitoT remained stable during CCCP treatment (Fig. 3H), evidencing that depolarisation-induced degradation is not universal to membrane tethering proteins. Thus, the depolarization-activated degradation might be specific for proteins arrested in the TOM translocase.

### Proteasome and VCP/p97 are not involved in the initial proteolytic clearance of the clogging fusion

In yeast *S. cerevisiae*, the ubiquitin-dependent chaperone Cdc48 is recruited to clogged TOM translocases to catalyze stalled cargo extraction. The extracted protein is subsequently directed for processing by the proteasome (Martensson et al., 2019). To test if an analogous mechanism is involved in the clearance of clogging fusion in our human model, we have targeted VCP/p97, a human ortholog of Cdc48, using its cell-permeable inhibitor NMS-873. VCP/p97 is an integral component of ubiquitinated protein degradation. Thus, its inhibition led to a general accumulation of ubiquitinated species in the cells (Fig. 4A). VCP/P97 inhibition showed no effect on the levels of full-size clogging fusion, but it increased the accumulation of processing forms of the fusion. Increased accumulation of the fusion fragments indicates that these are VCP substrates, which is likely due to their cytosolic localization. Notably, the NMS-873 inhibitor did not affect depolarisation-induced degradation. The full-size protein was also effectively cleared when VCP/p97 was inhibited.

**Fig. 4.**
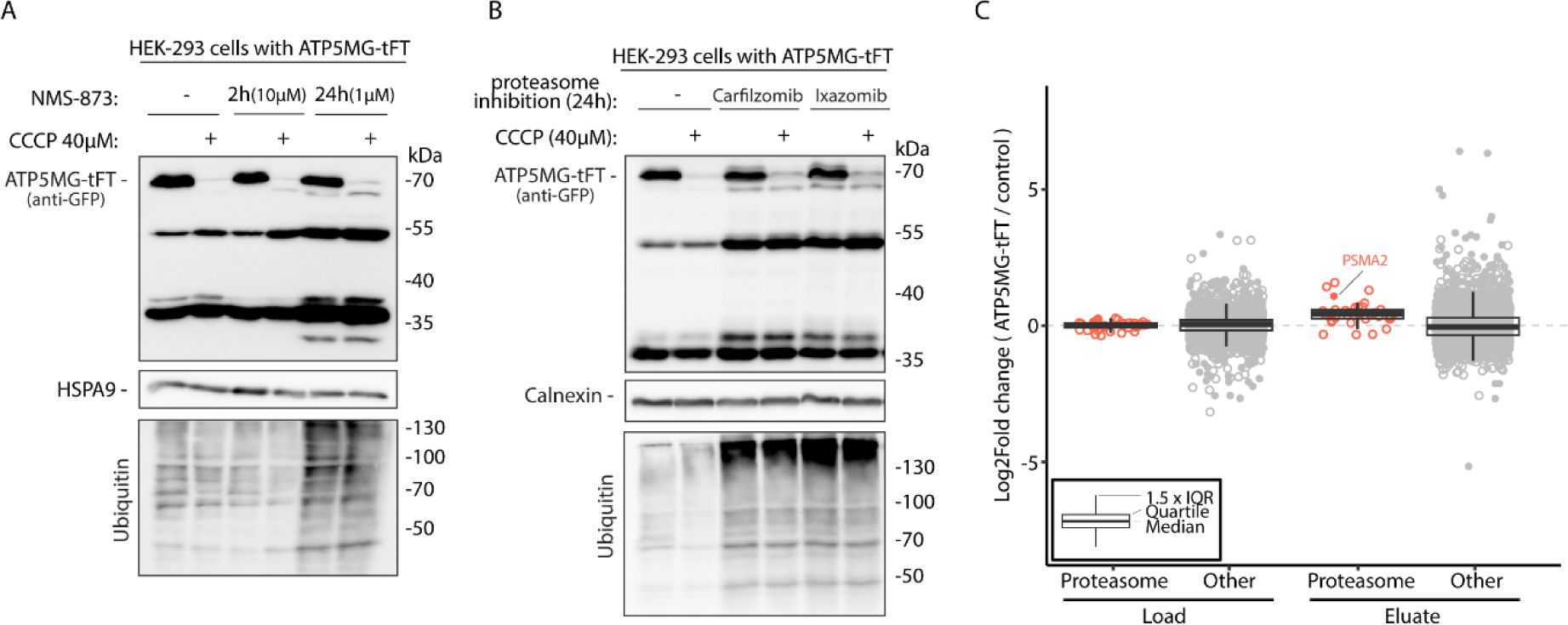
Mitochondrial depolarization-triggered processing of the stalled protein is not proteasome- or VCP/P97-dependent. (A, B) HEK-293 cells expressing ATP5MG-tFT (24 h tet. induction) were treated with indicated inhibitors: (A) P97 inhibitor NMS-873 was applied as indicated; (B) Proteasome inhibitors Carfilzomib and Ixazomib were applied for 24 h, parallel to induction; CCCP was added for 2 h. Cell extracts were analyzed by SDS-PAGE, western blotting, and tested with anti-GFP and control antibodies as indicated. (C) Jitter plot of protein log_2_ fold change (measured by LC-MS) in lysates (Load) or IP eluates from HEK-293 cells with ATP5MG-tFT vs. control cells not expressing the fusion (data available in Table S1). Proteasome components and other identified proteins are represented in red and grey, respectively. Filled circles indicate significant changes (Student’s t-test p<0.05, |log_2_ fold change| >1).

We next investigated how the proteasome impacts ATP5MG-tFT. We treated the cells with proteasome inhibitors Carfilzomib and Ixazomib. The inhibition was effective, as evidenced by the cellular accumulation of ubiquitinated species (Fig. 4B). Again, however, we observed no significant influence on the accumulation of full-size clogging protein. Mitochondria uncoupling still effectively cleared the translocase-blocking protein despite the proteasome being inhibited. Fragments of the fusion protein appeared to be substrates of proteasomal degradation as they were stabilized similarly to VCP inhibition.

In addition, we examined proteomic data from Fig. 2B for the status of proteasome components. The abundance of proteasome subunits remained unchanged in the cells expressing ATP5MG-tFT as compared to control cells (Fig. 4C, see Load). Thus, the expression of clogging fusion in the tested conditions did not result in stress-induced proteasome upregulation, contrary to what was observed in yeast (Boos et al., 2019).

In the co-IP with GFP-Trap, proteasome subunits appear to be slightly enriched with ATP5MG-tFT. Although only PSMA2, an alpha subunit of the 20S core, passed the significance criteria (Fig. 4C, see Eluate). The slight enrichment could result from the proteasome involvement in degrading fusion fragments that are present in the cytosol. Such fragments were copurified with GFP-trap beads in parallel to the complete clogging protein (see Fig. 2A).

In tandem with the ubiquitin-proteasome system, bulk degradation by autophagy provides clearance of cellular proteins. We observed the lysosome probe Lysotracker to localize in proximity to the fusion protein’s fluorescent signal only in a few cases (Fig. S3A). Moreover, treatment with Bafilomycin A, a lysosomal proteolysis inhibitor, had no apparent impact on the accumulation of ATP5MG-tFT protein (Fig. S3B). Bafilomycin A did not alter the fusion protein clearance upon depolarization. Finally, we did not observe any changes to the lysosome compartment as assayed with Lysotracker staining when comparing the cells with or without fusion induction (Fig. S3C). We thus conclude that autophagy did not significantly contribute to the translocase clogging response in our model.

Besides bulk protein disposal, autophagy also provides a selective protein degradation termed chaperone-assisted selective autophagy (CASA). HSP70 chaperones are a key component of CASA, working with co-chaperones and ubiquitin ligases to bring misfolded cargo to the autophagosome formation site. Stalled precursor proteins cannot fold properly and might expose hydrophobic fragments for HSP70 recognition. Thus we have treated the clogged cells with the HSP70 inhibitor VER-155008 (Massey et al., 2010). This treatment did not oppose CCCP-induced processing, indicating that HSP70 is not required for this process (Fig. S3D). Still, we observed a general reduction in the fusion levels. As both full-size and degradation products were decreased, we attribute this change to decreased fusion production and not increased degradation.

### Mitochondria-dependent processing of the clogging fusion

Our observations up to this point indicated that the initial proteolytic processing of the clogging fusion does not depend on the cytosolic QC machinery. This process is regulated by mitochondrial membrane polarization and thus possibly mediated by internal proteases. Only following this internal processing, fragments of the clogging protein would then be further degraded outside the organelle.

To verify this internal processing hypothesis, we developed an assay in which we monitored the state of the clogging protein in isolated mitochondria incubated in conditions differentially affecting the membrane potential (Fig. 5A). We used: (i) a trehalose-based isolation buffer, which assures correct osmotic conditions but does not contain metabolic substrates necessary to maintain mitochondrial membrane potential during prolonged incubation (Gnaiger et al., 2000; Pesta and Gnaiger, 2012; Hattori et al., 2005), and (ii) high-resolution respirometry buffer MIR05, supplemented with ADP+Mg^2+^, pyruvate, malate, and glutamate. The presence of respiratory substrates in the MIR05 allowed for keeping the mitochondria polarized during incubation. After one-hour incubation of isolated mitochondria at 37°C in the isolation trehalose buffer, the amount of clogging fusion in mitochondria was strongly decreased (Fig. 5B). In the case of mitochondria incubated in MiR05 buffer, the ATP5MG-tFT fusion remained largely unaffected.

**Fig. 5.**
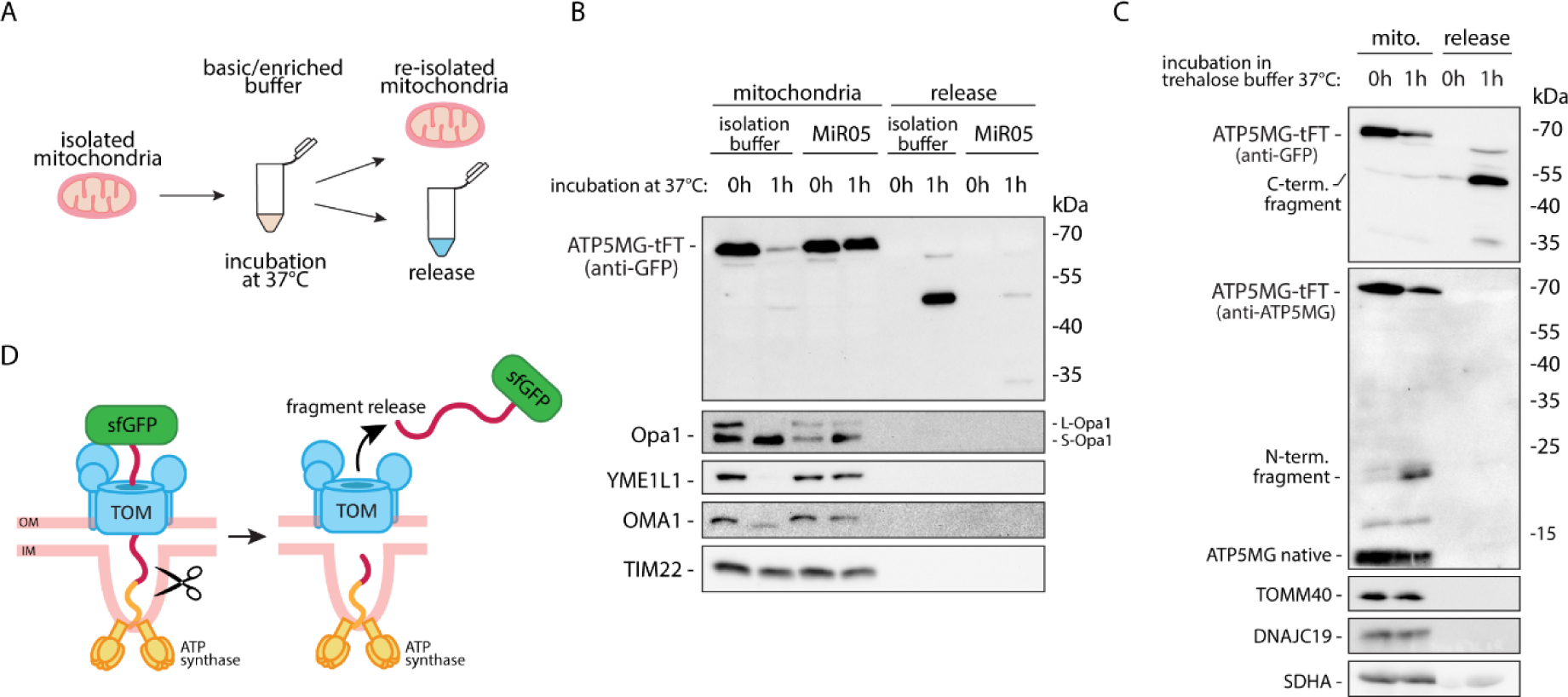
Isolated mitochondria process and release stalled protein when deprived of metabolic substrates. (A) Experimental scheme for B and C. (B, C) Mitochondria isolated from HEK-293 cells expressing ATP5MG-tFT were incubated in the isolation buffer or Mitochondrial Respiration Medium (MiR05) buffer supplemented with respiration substrates for the indicated time at 37°C. Next, mitochondria were separated from the buffers by centrifugation. Both fractions were analyzed by SDS-PAGE, western blot, and tested with anti-GFP and antibodies for mitochondrial proteins, as indicated. Anti ATP5MG antibody was used in (C) to visualize N-terminal part of the fusion. (D) Schematic representation of the intramitochondrial cleavage of the protein stalled in the TOM translocase.

The loss of ATP5MG-tFT protein from mitochondria upon incubation in trehalose buffer was accompanied by the processing of proteases OMA1 and YME1L1 and their substrate Opa1. These are hallmarks of mitochondrial response to stress, including depolarization (Baker et al., 2014; Zhang et al., 2014). Mitochondria incubated in MiR05 buffer, where ATP5MG-tFT was preserved, did not display pronounced processing of stress markers.

Importantly, the depletion of full-size clogging protein from mitochondria under stress was accompanied by the recovery of its smaller fragments in the release fraction (i.e., precipitated from the incubation buffer; Fig. 5B, see release). This indicates that stress-activated proteolytic cleavage allows the release of the cargo stalled in the OM translocase.

The anti-GFP antibody decorates the C-terminal part of the fusion protein (Fig. 1A). To test the fate of its N-terminal part, we used ATP5MG-specific antibodies (Fig. 5C). Both antibodies detected the full-size clogging fusion, which becomes depleted upon mitochondrial stress. However, contrary to the C-terminal fragment which was released from the organelle, the N-terminal part was retained in mitochondria with ATP5MG likely built into the IM.

Effective clearance of the clogging protein from isolated mitochondria indicates that the process does not require the recruitment of external factors. The necessary factors are present within the mitochondrial fraction and become activated by IM depolarisation. Notably, the model clogging protein was not cleared as a whole following proteolytic cleavage. The part arrested in the TOM translocase was released from the organelle, while the part integral to the IM remained inside mitochondria (Fig. 5 D).

### Mitochondrial metallopeptidase OMA1 is involved in the depolarization-dependent degradation of the clogging protein

Many mitochondrial proteases contain metal ions in their catalytic centers. We have treated the cells with metal ion chelator *o*-phenanthroline to inhibit the functions of metalloproteases. This treatment largely prevented CCCP-induced degradation of the clogger (Fig. 6A). The chelating agent was also effective in organello, limiting ATP5MG-tFT processing upon mitochondrial stress resulting from the incubation in the trehalose-based isolation buffer (Fig. 6B). These results suggest that the uncoupling-induced processing of clogged proteins, at least partly, depends on metalloprotease activity.

**Fig. 6.**
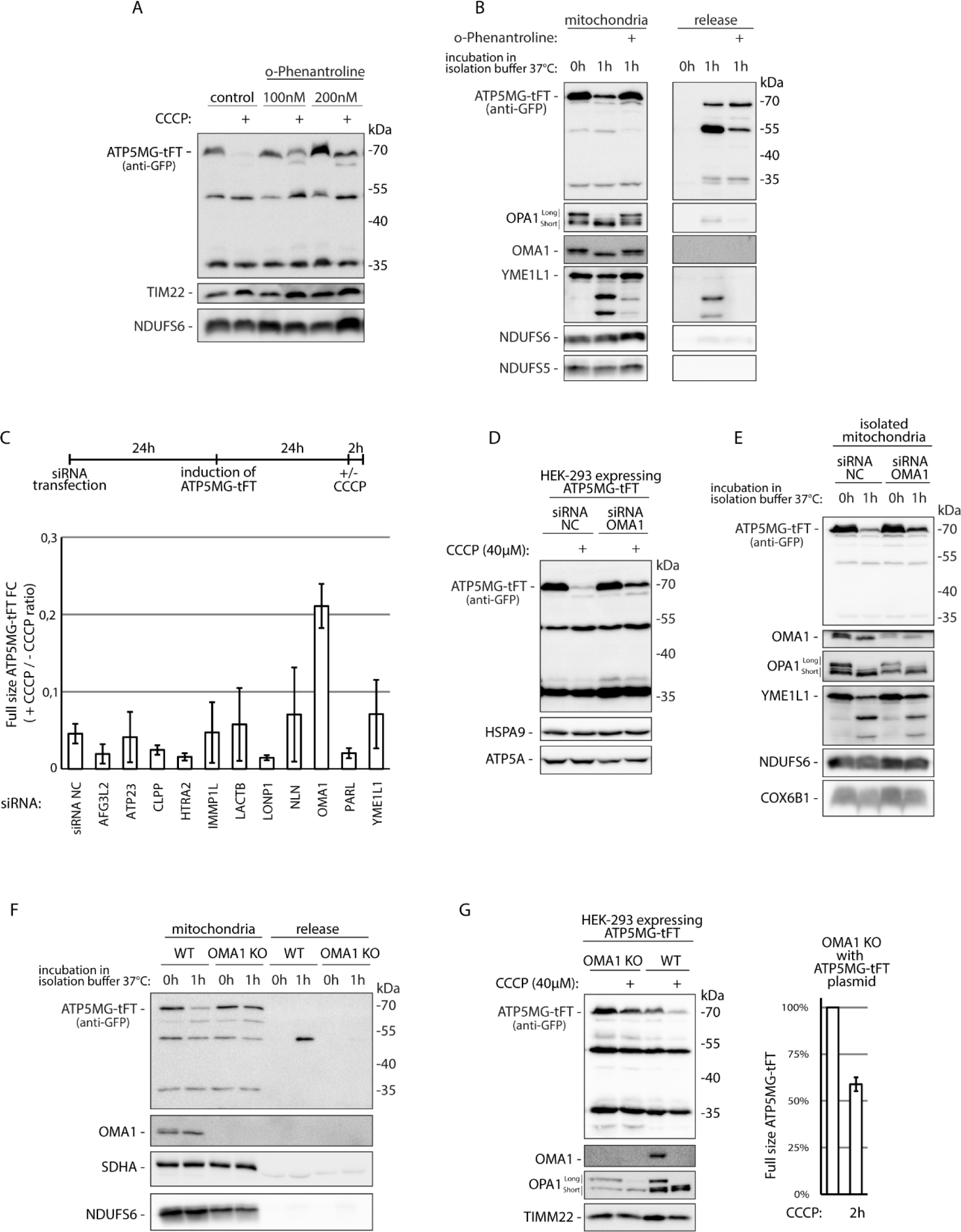
Mitochondrial metallopeptidase OMA1 contributes to depolarization-dependent degradation of the mitochondrial protein translocase stalled cargo. (A) HEK-293 cells with ATP5MG-tFT were treated with CCCP, in the presence or absence of *o*-phenanthroline, for 2h. (B) Isolated mitochondria with ATP5MG-tFT were incubated in isolation buffer with or without *o*-phenanthroline for indicated time at 37°C. Mitochondria and release fractions were analysed by western blot. (C) HEK-293 cells with ATP5MG-tFT were treated with siRNA targeting selected mitochondrial proteases. Levels of the fusion protein were tested by western blotting and quantified by densitometry. Graph indicates fold change in the levels of full size fusion protein during 2 h of CCCP treatment. Mean values +/- SEM from n=3 independent experiments are presented. (D) HEK-293 cells expressing ATP5MG-tFT were treated with OMA1 targeting or control siRNAs for 72h, induced with tetracycline for 24 h; CCCP treatment was applied for 2h. (E) Mitochondria isolated from HEK-293 cells expressing ATP5MG-tFT and treated with OMA1 targeting or control siRNAs for 48 h, were incubated in isolation buffer for indicated time at 37°C. Re-isolated mitochondria were analysed by western blotting with indicated antibodies. NC - negative control. (F) Mitochondria isolated from WT or OMA KO HEK-293 cells transfected with ATP5MG-tFT expressing plasmid were incubated in isolation buffer for indicated time at 37°C. Re-isolated mitochondria and coresponding release fractions were analyzed by western blotting with indicated antibodies. (G) HEK-293 OMA1 KO and HEK-293 cells transfected with plasmid encoding ATP5MG-tFT were treated with or without CCCP for 2h. Levels of the fusion protein were tested by western blotting. Graph indicates the levels of full size fusion protein following CCCP treatment; densitometry values for untreated cells were set to 100%. Mean values +/- SEM from n=4 independent experiments are presented.

We targeted an array of mitochondrial proteases with siRNA to test for their potential role in clogging fusion degradation. Following cell treatment with specific siRNA, we have compared the levels of full-size ATP5MG-tFT fusion in standard conditions, and after CCCP treatment (Fig. 6C). Out of the tested siRNA, only the one targeting OMA1 metalloprotease partially prevented the CCCP triggered processing. Silencing OMA1 decreased the degradation of arrested protein in CCCP-treated cells (Fig. 6D) and isolated mitochondria (Fig. 6E). However, the silencing effect was only partial. This could be a consequence of the incomplete depletion of OMA1 (see Fig. 6E blot for OMA1), or of another protease contributing to the process. To further investigate the role of OMA1, we turned to a knockout cell line (OMA1 KO) (Baker et al., 2014). We compared mitochondria isolated from wild-type and OMA1 KO HEK-293 cells following transient transfection with ATP5MG-tFT expressing plasmid (Fig. 6F). Upon incubation in stress-inducing conditions, ATP5MG-tFT fusion was largely protected from cleavage and release in organelles lacking OMA1 as compared to these isolated from unmodified cells. Accordingly, in a cell-based assay, we observed increased stability of the full-size clogging fusion in the absence of OMA1 protease (Fig. 6G). However, even in OMA1 KO cells, ATP5MG-tFT fusion was destabilized upon treatment with the uncoupling agent. The western blot signal decreased to 59% of the signal before adding any uncoupling agent (CCCP, Fig. 6G see graph). This finding indicates that while OMA1 is a major processor, the depolarisation-triggered processing of protein arrested in transit is likely not restricted to OMA1 activity and could be executed by another protease.

### Stalled import intermediates disturb cristae morphology

Our observation that ATP5MG-tFT can physically bind OM-located TOM complex and IM- located ATP synthase (Fig. 2) drove us to investigate its impact on mitochondrial ultrastructure. A protein stalled in transit and spanning two membranes is likely to impact mitochondrial morphology. This is especially relevant since ATP synthase complexes are distal to the OM as they locate in crista membranes often assembled into rows of dimers along the curved crista lamellae edges (Strauss et al., 2008; Rampelt et al., 2022).

Using transmission electron microscopy (TEM), we compared the architecture of mitochondrial membranes in the cells with or without expression of ATP5MG-tFT (Fig. 7A,B). We found that the clogging protein strongly altered mitochondrial morphology. In cells expressing the clogging construct, most of the mitochondria had a significantly reduced number of cristae (69.9 % of mitochondria with abnormal cristae; Fig. 7B, abnormal type A and B). The remaining crista membranes were often positioned along organelles’ perimeters resulting in a void appearance of the matrix. In some cases, crista lamellae formed elongated round structures. At the same time, the control cells not expressing ATP5MG-tFT predominantly contained normal mitochondria (89.2 %).

**Fig. 7.**
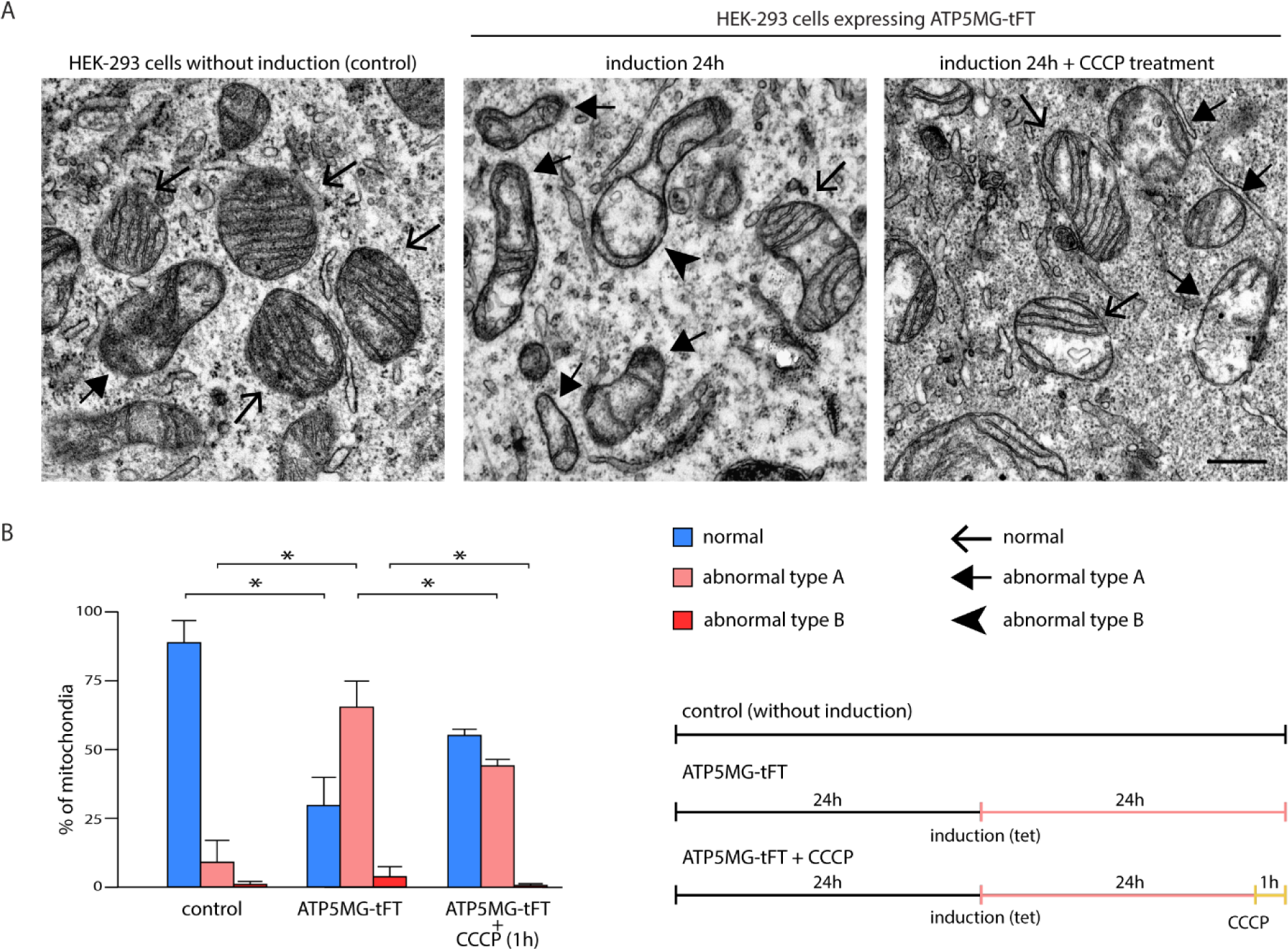
Disturbance of cristae morphology caused by stalled import intermediates is reversed by mitochondria uncoupling. (A) Transmission electron microscopic imaging of mitochondrial ultrastructure in HEK-293 cells with or without ATP5MG-tFT expression and with or without CCCP treatment. Representative images are shown. Scale bar 500 nm. (B) The percentage of mitochondria with normal or aberrant cristae was quantified. Mean + error bars (SEM), n=4 biological replicates with 8-11 profiles per replicate; Treatments were compared by Mann-Whitney U test; *p<0.05. Experimental scheme is shown at the bottom right.

As uncoupling-induced cleavage releases the fusion protein, it should effectively disengage the two mitochondrial membranes. To test this possibility, we examined the effect of CCCP treatment on mitochondrial ultrastructure in cells expressing clogging protein (Fig. 7B). The uncoupler treatment is known to disturb mitochondrial morphology and reduce the number of normal cristae (Viana et al., 2021; Miyazono et al., 2018). However, we observed that uncoupling-induced clogger release resulted in significant restoration of the cristae structure, seen in TEM (during one hour of CCCP treatment, the fraction of mitochondria with normal cristae increased from 30.1 % to 55.2 %; the fraction of mitochondria with abnormal cristae decreased from 69.9 % to 44.8 %). This observation supports the assumption that arrested precursors disrupt mitochondrial structure by clasping OM and cristae membranes. Moreover, this proves that clogging-induced damage is not permanent and can be reversed by the mitochondrial QC machinery.

## Discussion

Import of proteins into mitochondria gains increasing recognition as a critical component of the cellular proteostasis network. The undisturbed flow of incoming protein precursors is essential to maintain organellar functions and prevent protein mislocalization. However, the flow can be interrupted if proteins stall in transit and block translocases. We and others provide clear evidence of the detrimental consequences of import disturbance for mitochondrial and cellular physiology. Thus, protein translocation failure requires adequate quality control (QC) responses. Cellular responses to stress caused by disturbed protein import have primarily been defined in the yeast model (Lenkiewicz et al., 2022). Fusion proteins designed to stall during the import proved to be an effective discovery platform to characterize the molecular mechanisms that resolve translocation failure in yeast (Boos et al., 2019; Martensson et al., 2019; Weidberg and Amon, 2018). Using a similar strategy, we created an import arrest model in a cultured cell line to reveal mechanisms of import QC in human cells and provide their detailed description. We have fused ATP5MG protein, a F*_O_* ATP synthase component, with a tFT tag to cause precursor protein stalling in mitochondrial translocases. The stably folding sfGFP, a part of tFT, is not translocated effectively (Gomkale et al., 2021; Kowalski et al., 2018). Our results confirmed the translocation arrest of ATP5MG-tFT. The fusion localized to mitochondria, but its C-terminus remained exposed on the organelle’s outside, indicating incomplete import. Proteomic analysis revealed a pronounced interaction of the fusion not only with TOM translocase but also with the ATP synthase complexes, supporting IM integration of its N-terminal part. As expected in translocase blockage, model fusion protein interfered with the import of native mitochondrial proteins, impaired mitochondrial function, and reduced cellular proliferation.

Yeast cells respond to import disturbance by broadly adapting their proteostasis network. Such changes, executed at the transcriptional and post-transcriptional levels, modulate protein turnover by adjusting protein synthesis and degradation rates. Among post-transcriptional changes, disturbed protein import inhibits protein synthesis and simultaneously boosts the activity of the proteasome in the processes called UPRam (unfolded protein response activated by mistargeting of proteins)(Topf et al., 2016) or mPOS (mitochondrial precursor overaccumulation stress)(Wang and Chen, 2015). Proteasome upregulation during mitochondrial dysfunctions, which affect protein import, is also mediated transcriptionally and is not limited to yeast (Sladowska et al., 2021; Kim et al., 2021; Boos et al., 2019).

Individual mitochondrial stresses can differentially alter the transcriptome but frequently trigger the integrated stress response (ISR) (Quirós et al., 2017; Topf et al., 2019; Kaspar et al., 2021; Fessler et al., 2020; Guo et al., 2020). ISR reduces overall protein synthesis while allowing the translation of some specific ORFs, followed by transcriptome adjustment. ISR-induced changes alleviate import stress by decreasing the synthesis of mitochondria-targeted precursor proteins and stimulating general proteostasis restoration. However, more direct mechanisms are required to unblock translocases. In yeast, a mechanism related to endoplasmic reticulum-associated degradation (ERAD) operates at the OM to remove stalled precursors. This pathway, named the mitochondrial protein translocation-associated degradation (mitoTAD), acts at the TOM complex (Martensson et al., 2019). MitoTAD involves Ubx2 protein recruting cytosolic AAA-ATPase Cdc48 to extract the arrested precursor protein and transfers them for proteasomal clearance. The Rsp5 ubiquitinates precursors assisting their degradation (Schulte et al., 2023). A parallel QC pathway that controls unimported precursors uses Pth2 protein, which binds to TOM independently of Ubx2. Pth2 recruits the cytosolic ubiquilin family protein Dsk2, which assists in the proteasomal degradation of ubiquitinated precursors (Schulte et al., 2023). However, it is not fully clear if Pth2 acts on precursors that stall in the translocase channels or those that bind translocase receptors.

In parallel to constitutive QC, inducible mechanisms were shown to respond to import arrest in yeast. Such responses are induced once import-related stress exceeds the capacity of basic ones and are also based on stalled protein extraction for proteasomal processing. One such mechanism, termed mitochondrial compromised protein import response (mitoCPR), activates upon precursors overload of the TIM23 translocase (Weidberg and Amon, 2018). In such conditions, transcription factor Pdr3 induces Cis1 protein expression. Cis1 localizes at the OM and recruits mitochondrial AAA-ATPase Msp1 to the clogging site. Msp1 extracts the arrested protein allowing its further degradation by the proteasome.

The QC mechanism we report here fundamentally differs from those described in yeast cells. Degradation of translocase clogging proteins in yeast depends on active extraction by AAA ATPases coupled with degradation by the ubiquitin-proteasome system. The mechanism we uncovered using the human cell translocation arrest model is based on proteolytic cleavage of the arrested protein allowing back movement and release of its blocked portion from the TOM translocase. We observed that the arrested protein remains relatively stable and is cleared effectively only after depolarizing the IM. We provide evidence that the cleavage of arrested protein is mediated by mitochondrial proteases, likely within its fragment exposed to the IMS. Mitochondria have multiple proteolytic enzymes that regulate the organelle’s proteome and respond to protein damage (Szczepanowska and Trifunovic, 2021; Quirós et al., 2015). We confirmed the involvement of OMA1 metalloprotease in arrested precursor processing. OMA1 is characterized by a low basal activity until it is activated by stress, such as mitochondrial membrane depolarization (Baker et al., 2014; Zhang et al., 2014). The adaptable activity of OMA1 corresponds to the observed low basal degradation of the clogging fusion protein, amplified by ΔΨ dissipation. The proximity interactome of OMA1 included TOM components, further supporting this protease’s role in translocation QC (Botham et al., 2019). Still, the stabilization of the translocase-arrested protein was only partial in the absence of OMA1, suggesting that also another enzyme contributes to the arrested protein cleavage. Mitochondrial proteases share part of their substrates (Botham et al., 2019). A well-established example is the OMA1 and YME1L proteases pair, mediating the processing of OPA1 in the membrane potential-dependent manner (Rainbolt et al., 2016). A recent study reports a yeast homolog of YME1L - Yme1 to clear mutant forms of Aac1 protein, which can stall during import (Coyne et al., 2023). Moreover, the activity of several other proteases, like PARL, AFG3L2, LONP1, or ClpXP, is also regulated by the membrane potential (Sekine et al., 2012; Pryde et al., 2016; Patron et al., 2022). Still, our siRNA-based silencing experiments did not support the involvement of any of these proteases in the degradation of the stalled cargo. Transient silencing might not sufficiently reduce targeted enzyme levels, which is a limitation of the method. It is also conceivable that OMA1 acts indirectly by activating another protease to mediate the release. However, the redundancy of proteases appears to be a more likely interpretation ensuring the robustness of the QC machinery that controls the import.

Remarkably, the role of proteasomal degradation appears indirect and more limited in comparison to QC mechanisms discovered in yeast. We did not observe increased proteasome subunits levels in cells expressing ATP5MG-tFT. We also did not find the proteasome among significantly enriched partners of the fusion protein. Still, ATP5MG-tFT underwent degradation even without uncoupler treatment, as manifested by the accumulation of smaller protein fragments. Such fragments are characteristic of tFT fusions and represent degradation intermediates (Khmelinskii et al., 2016). In contrast to the whole fusion protein, these fragments were significantly stabilized in response to the inhibition of the proteasome or VCP/p97, a human homolog of yeast Cdc48. At the same time, the depolarisation-triggered processing of the full-size protein was unaffected by the inhibitors. Processing intermediates were found outside mitochondria, but originated from the mitochondrially localized full-size ATP5MG-tFT fusion. Notably, their release did not require the recruitment of external factors, a fundamental difference in comparison to QC mechanisms identified in yeast.

The QC response mediated by internal components of mitochondria appears largely independent of cytosolic translation modulation. Likewise, in the time frame of our experiments, we did not observe substantial proteome remodeling that would indicate ISR activation. We also did not detect the activation of alternative stress-response programs, like the UPRmt, with no changes in CLPP or heat-shock proteins (Fessler et al., 2020). Still, we cannot entirely exclude ISR activation as only a subset of known targets was detected in our proteomic dataset (Supplementary Table 1) (Neill and Masson, 2023). In classic mitochondrial genetics, a threshold effect is known, with cells tolerating high levels of mutated mtDNA while maintaining proper function and sufficient respiratory capacity (Wallace and Chalkia, 2013). However, once the mutation load exceeds the tolerated threshold, mitochondrial functions collapse. The induction of ATP5MG-tFT expression only mildly affected cell proliferation, which may reflect its moderate expression. Assuming that a fraction of translocation events continually fail under physiological conditions, no impact at the cellular level may be detectable due to their number staying below a threshold.

Depolarisation triggered QC not only cleared the arrested protein but also allowed the restoration of the architecture of mitochondrial membranes. Stalled precursor proteins can form tethers connecting mitochondrial membranes. If such tethers are directed to cristae, the lamellar invaginations of IM, their limited length can distort the membrane’s architecture. This appears to be the case for the model protein we used. The expression of ATP5MG-tFT resulted in a significantly reduced number of cristae and their location mainly alongside the inner boundary membrane. Cristae morphology is pivotal for respiratory chain function. Consequently, arrested proteins can impair respiratory chain function and, without effective QC, would propel further import stalling. Remarkably, the uncoupler-induced clearance of the arrested ATP5MG-tFT restored the cristae architecture. Mitochondrial proteases, including OMA1, are among the regulators of organellar ultrastructure. OMA1 KO mitochondria display strongly disturbed morphology (Viana et al. 2021), resembling the phenotype we observed upon ATP5MG-tFT expression. OMA1 is a well-established regulator of OPA1 dynamin-like GTPase, a part of the machinery shaping IM (Quintana-Cabrera and Scorrano, 2023). Though, given the role of OMA1 in releasing arrested proteins, the tethering of mitochondrial membranes by stalled import intermediates should be considered among mitochondria shaping factors.

In our study, we fused mitochondrial protein with a tag to model stalling in translocation. This approach proved effective but might be regarded as synthetic. However, multiple data substantiate that the phenomenon we modeled also affects native mitochondrial proteins that can become arrested upon import, particularly when expressed in excess or mutated (Weidberg and Amon, 2018; Boos et al., 2019; Coyne et al., 2023). Moreover, mitochondrial translocases were shown to engage with misfolded or aggregated proteins from the cytosol even if such proteins were not destined for mitochondria (Ruan et al., 2017). Such events were observed in neurodegenerative disorders associated with protein misfolding, including Alzheimer’s disease (AD) and Parkinson’s disease (PD). In AD brain samples, amyloid precursor protein (APP) was found to interact with TOMM40 and TIMM23 in mitochondria (Devi et al., 2006). Moreover, APP expressed in cultured cells stalled in translocases, disturbing mitochondrial protein import (Anandatheerthavarada et al., 2003). APP-derived amyloid-β peptides were also found to suppress mitochondrial precursor processing, increasing the levels of these immature protein forms (Mossmann et al., 2014). Likewise, the PD-associated variant of α-synuclein was observed to associate with mitochondria, interact with TOM translocase, and inhibit mitochondrial protein import (Gallardo et al., 2008; Di Maio et al., 2016). Similarly, huntingtin, another disease-linked protein, was proposed to block mitochondrial translocases (Yano et al., 2014). The above examples of clinically relevant proteins substantiate the risk of clogging for mitochondrial protein translocases. In a possible link of these observations to our discoveries, the uncoupling agent DNP was found to improve functioning in animal models of neurodegenerative diseases related to protein misfolding (Wu et al., 2017). Partial depolarization of mitochondria was recently also suggested as part of the cellular program that counteracts aging-related decline (Vyssokikh et al., 2020). It is an intriguing possibility that drugs altering mitochondrial membrane potential, generally regarded as protein import inhibitors, could help in import restoration. We thus propose that applying models of clogging is necessary to streamline the discovery and understanding of protein import QC in mammalian cells.

Our research leaves open points which require addressing. Following the cleavage, part of the arrested protein remains inside mitochondria. Such protein fragments could have deleterious effects and likely are further processed by mitochondria. The released fragments appear to fall under the general control of the ubiquitin-proteasome system. However, their impact on proteostasis should not be overlooked. Recent research proves that unimported mitochondrial proteins can aggregate disturbing proteostasis (Nowicka et al., 2021). Hence, cytosolic chaperones might be necessary to prevent their toxicity (Krämer et al., 2023). The cytosolic handling of mitochondria-released fragments is a significant part of their QC requiring further attention.

## Materials and methods

### Cell lines and growth conditions

HEK-293, HEK-293 Flp-In T-REx, and HeLa cells were cultured at 37°C with 5% CO_2_ in standard Dulbecco’s modified Eagle’s medium (DMEM) containing 4500 mg/L glucose, supplemented with 10% (v/v) fetal bovine serum, 2 mM l-glutamine, 100 U/ml penicillin, and 100 μg/ml streptomycin sulfate. Flp-In T-REx HEK-293 with inducible expression of ATP5MG-tFT or IleDeg-tFT were generated by co-transfection with pcDNA5/FRT/TO plasmid encoding respective fusion protein and a helper plasmid pOG44. Stable transgenic cells were selected with hygromycin treatment. The resulting cell lines were tested for inducible expression of tFT fusion by adding tetracycline (1 µg/ml) for 24 h as described previously (Chojnacka et al., 2022). OMA1 KO T-Rex HEK-293 cells were kindly provided by Thomas Langer (Baker et al., 2014). HEK-293 and HeLa were acquired from the American Type Culture Collection (ATCC; www.atcc.org). Cell proliferation was measured by direct cell counting using a cell counter (Countess II; Life Technologies) or manually with a hemocytometer.

### Total cellular protein lysate

Cells were harvested with trypsin and lysed in RIPA buffer (65 mM Tris–HCl [pH 7.4], 150 mM NaCl, 1% v/v NP 40, 0.25% sodium deoxycholate, 1 mM ethylenediaminetetraacetic acid [EDTA], and protease inhibitor cocktail (Merck, P8340)) for 30 min at 4°C. After 10 min centrifugation at 5 000 x g at 4°C, supernatants were collected, and the protein concentration was measured by Bradford assay (ROTI Quant, ROTH). Proteins of the lysate were precipitated by the addition of trichloroacetic acid (TCA) to the concentration of 12.5%. and pelleted by centrifugation at 20 000 x g at 4°C. The pellets were washed with acetone, followed by solubilization and denaturation with Urea Sample Buffer (6M urea, 6% SDS (288.38g/mol), 125mM Tris-HCl, pH 6.8, 0.01% Bromophenol Blue, and 50 mM DTT) at 65°C.

### Cell transfection

Cells were transfected with plasmid DNA using Gene Juice Transfection Reagent (Merck Millipore) 24 h after plating. Transfection was performed according to the manufacturer’s protocol. Cells were harvested 24-48 h post-transfection as indicated.

### Cell fractionation

Cells were harvested with trypsin and resuspended in Isolation buffer (IB; 20mM Hepes pH 7.6, 220 mM mannitol, 70 mM sucrose, 1 mM EDTA, 2 mM PMSF). Cells were homogenized in a Dounce glass homogenizer, and the homogenate was subjected to centrifugation at 1 000 x g for 10 min at 4°C to pellet the cellular debris. The supernatant was carefully removed. Part of it was stored as a total fraction, and the rest was subjected to subsequent centrifugation at 10 000 x g for 10 min at 4°C. After centrifugation, supernatant containing cytosolic proteins and mitochondria-enriched pellets were separated. To test for protease accessibility, mitochondrial samples were treated with proteinase K (25μg/ml) for 15min on ice. Proteinase K was inhibited by 2mM PMSF followed by pelleting mitochondria at 20 000 x g for 10 min at 4°C. Aqueous fractions (i.e., total and cytosol) were precipitated with TCA as described for total cell extracts. Pelleted proteins or mitochondria were solubilized and denatured with Urea Sample Buffer, and analyzed by western blotting. For nuclear fraction, Abcam Cell Fractionation Kit (ab109719-1) was used, in accordance with the manufacturer’s recommendations.

### Mitochondrial isolation and model protein processing assay

Mitochondria isolation was performed as described previously (Mohanraj et al., 2019). Briefly, cells were harvested, resuspended in ice-cold isolation trehalose buffer (300 mM trehalose, 10 mM HEPES-KOH pH 7.7, 10 mM KCl, and 2 mg/ml of bovine serum albumin (BSA)), and homogenized in a Dounce homogenizer. Homogenates were clarified by centrifugation at 650 x g at 4°C. The resulting supernatant was centrifuged at 14 000 x g for 15 min at 4°C to obtain a mitochondrial pellet. Mitochondria enriched pellet was resuspended in isolation trehalose buffer without BSA or MiR05 Mitochondrial Respiration Medium (Gnaiger et al., 2000), as indicated. Mitochondria were used directly without storage. Samples were incubated at 37°C for 1 h or analyzed without incubation to observe the processing and release of the model protein. Mitochondria were re-isolated by centrifugation, and the supernatant was collected as a release fraction. Release fractions were precipitated by the addition of TCA as described for total cell extracts. Both mitochondria and release samples were denatured with the Urea Sample Buffer and analyzed by western blotting.

### siRNA mediated knockdown

Cells were reverse transfected with siRNA targeting selected genes (MISSION esiRNA; Merck) or non-targeting siRNA control (MISSION siRNA universal negative control UNC2; Merck) using Lipofectamine RNAiMAX Reagent (ThermoFisher). For each well of a 6-well plate, 3μl of Lipofectamine RNAiMAX Reagent (ThermoFisher) was diluted in 250μl of Opti-MEM reduced serum medium (Gibco) and siRNA was diluted in 250μl Opti-MEM medium. Subsequently, both were mixed inside the well to prepare complexes. After 5 min of incubation, 2 ml of cells suspension in growth media were added. Cells were harvested after 48h or 72h, as indicated. The final concentration of siRNA was 20nM.

### LC-MS/MS analysis and data processing

Chromatographic separation was performed on an Easy-Spray Acclaim PepMap column 50 cm × 75 µm inner diameter (Thermo Fisher Scientific) at 45°C by applying 135 min acetonitrile gradients in 0.1% aqueous formic acid at a flow rate of 300 nl/min. An UltiMate 3000 nano-LC system was coupled to a Q Exactive HF-X mass spectrometer via an easy-spray source (all Thermo Fisher Scientific). The Q Exactive HF-X was operated in data-dependent mode with survey scans acquired at a resolution of 60,000 at m/z 200. Up to 12 of the most abundant isotope patterns with charges z=2-6 from the survey scan were selected with an isolation window of 1.3 m/z and fragmented by higher-energy collision dissociation (HCD) with a normalized collision energy of 27, while the dynamic exclusion was set to 30 s. The maximum ion injection times for the survey scan and the MS/MS scans (acquired with a resolution of 15,000 at m/z 200) were 45 and 22 ms, respectively. The ion target value for MS was set to 3e6 and for MS/MS to 1e5, and the intensity threshold for MS/MS was set to 8e2. The obtained data were processed with MaxQuant v. 1.6.7. 0 (Cox and Mann, 2008), and the peptides were identified from the MS/MS spectra searched against Human Reference Proteome UP000005640 using the build-in Andromeda search engine. Raw files corresponding to 12 affinity purified samples (3 conditions, 4 replicates) and 12 input samples (3 conditions, 4 replicates) were loaded into MaxQuant and processed. Cysteine carbamidomethylation was set as a fixed modification and methionine oxidation as well as protein N-terminal acetylation were set as variable modifications. For in silico digests of the reference proteome, cleavages of arginine or lysine followed by any amino acid were allowed (trypsin/P), and up to two missed cleavages were allowed. The False Discovery Rate (FDR) was set to 0.01 for peptides, proteins and sites. Match between runs and LFQ quantification were enabled. Other parameters were used as pre-set in the software. Unique and razor peptides were used for quantification enabling protein grouping (razor peptides are the peptides uniquely assigned to protein groups and not to individual proteins).

LFQ intensities for protein groups were loaded into Perseus v. 1.6.6.0 (Tyanova et al., 2016) and log2 transformed. Proteins that were only identified by site, reverse, or potential contaminant were removed. Protein groups with LFQ intensity values in less than 2 out of 4 ATP5MG-tFT eluate samples were removed. Further data analysis and visualizations were performed in R v. 4.2.1 using the libraries tidyverse v. 1.3.2, fuzzyjoin v. 0.1.6, ggrepel v. 0.9.1. Missing values were attributed to low abundances and imputed by drawing from a normal distribution with a mean shifted down by 1.8 standard deviations and a standard deviation narrowed by a factor of 0.3. Imputation was performed separately for each group (i.e., total cell lysate or IP) to account for different global compositions. Fold changes were calculated as the difference between each group’s log2 average value. P-values were calculated with a two-sided, paired Student’s t-test. No multiple comparison correction was applied. Statistical significance was assigned above an absolute fold change of 1 and p-value below 0.05 unless otherwise stated.

The raw mass spectrometry data have been deposited to the ProteomeXchange Consortium via the PRIDE (Perez-Riverol et al., 2022) partner repository with the dataset identifier PXD038583.

### Transmission electron microscopy (TEM)

HEK-293 cells (60%-70% of confluency) were fixed by 2.5% glutaraldehyde and 2% paraformaldehyde (both from Electron Microscopy Science, Inc.) solution for 1 h at 4°C. After fixation, cells were rinsed three times for 10 min with 0.1 M cacodylate buffer (BDH Chemicals Ltd.). Next, the cells were postfixed in 1% osmium tetroxide (Polsciences Inc.) for 1 h at room temperature, rinsed three times for 10 min with water, and stained with 2% uranyl acetate (Serva Electrophoresis GmbH) for 30 min. Dehydration was performed by incubating the sample in increasing acetone concentrations (25%, 50%, 70%, 90%, 96%, and 2 x 100%; each 10 min). After that, the cells were embedded in the mixture of pure acetone and Epon (Serva Electrophoresis GmbH) resin (1:1 and 1:2, each mixture for 30 min), then in pure Epon resin exchanged tree times (incubation: 1 h, overnight, and 1 h). After resin polymerization at 60°C, 60 nm thick sections were collected on TEM grids. Electron micrographs were obtained with Morada camera using a JEM 1400 transmission electron microscope at 80 kV (JEOL Co., Japan) in the Laboratory of Electron Microscopy, Nencki Institute of Experimental Biology of Polish Academy of Sciences, Warsaw, Poland.

### TEM Image analysis

TEM cellular studies were repeated four times for each experimental group (n=4). Quantification was obtained from 8-11 cell profiles for each repeated group. The number of mitochondria was established using Cell Counter plugin of ImageJ Fiji software (Schindelin et al., 2012), then the percentage of normal and abnormal mitochondria was determined. Results are presented as arithmetic means with standard errors. The significance of differences between the groups was tested with the Mann-Whitney U test using GraphPad Prism 8. The standard value of p <0.05 was adopted as the critical level of significance. The significance level was marked in the graphs as: * for p <0.05, ** for p <0.01, and *** for p <0.001. In case of no significance, the abbreviation “ns” was used.

### Immunofluorescent staining

Cells seeded on glass coverslips were fixed for 10 minutes at room temperature with 4% paraformaldehyde in PBS. Then they were washed with PBS three times for 5 min. Cell membranes were permeabilized with 0.1% Triton-X100 in PBS for 10 min, followed by three washes with PBS and blocking with 5% BSA in PBS for 1 hour at room temperature. Incubation with primary antibodies was performed overnight at 4⁰C in wet chamber. The antibodies were appropriately diluted in PBS with 2% BSA 1:50 for TOMM20 (Santa Cruz) and 1:1000 for calnexin (Abcam). After the incubation, three washes with PBS were applied and coverslips were incubated for 1h at room temperature with secondary antibodies diluted 1:1000 in 2% BSA in PBS. Then, three washes with PBS were applied once again and the coverslips were mounted on microscopic slides using ProLong Diamond Antifade Mountant with DAPI (Thermo Fisher Scientific). The stained cells were imaged using a confocal fluorescent microscope Leica SP8 with HC PL APO CS2 63x/1.4 oil objective.

### Life cell staining

Cells were incubated for 2 h at 37⁰C in a culture medium containing 50 nM Mito Tracker Deep Red (Thermo Fisher Scientific) and/or 75 nM LysoTracker Blue DND-22. Then, the stained cells were visualized using a confocal fluorescent microscope Leica SP8 with HC PL APO CS2 63x/1.2 water objective.

### High-resolution respirometry

High-resolution respirometry was performed as previously described (Chojnacka et al., 2022). Oxygen consumption was measured in intact cells using Oxygraph-2k. Data were digitally recorded using DatLab v. 5.1.0.20 (Oroboros Instruments, Innsbruck, Austria) and expressed as pmol of O_2_/min per 10^6^ cells following the manufacturer’s instruction for air calibration and background correction. Trypsinized cells were suspended at 1 to 2 × 10^6^ cells/mL in 2 mL of MiR05 medium (0.5 mM ethylene glycol-bis(β-aminoethyl ether)-N,N,N′,N′-tetraacetic acid [EGTA], 3 mM MgCl_2_, 60 mM lactobionic acid, 20 mM taurine, 10 mM KH_2_PO_4_, 20 mM HEPES, 110 mM d-sucrose, 1 mg/ml BSA–fatty acid-free added freshly, pH 7.1) and immediately placed into the Oxygraph chamber to access basal respiration. HEK-293 Flp-In T-REx derivative proved susceptible to blocking ATP-synthase (irrespective of ATP5MG-tFT expression). Concentrations of oligomycin A as low as 0.25 μM abrogated O_2_ consumption in these cells (data not shown). Therefore, we assessed the overall OCR cell profile without oligomycin A. The maximum respiration (electron transport system [ETS] capacity) was determined upon titration of the uncoupler CCCP (carbonyl cyanide m-chlorophenyl hydrazone) (0.20 μM/step). Next, complexes I, III, and IV were inhibited by rotenone (0.5 μM), antimycin A (5 μM), and sodium azide (50 μM), respectively, to detect residual oxygen consumption (ROX). Stable O_2_ flux plateaus were used for the calculation of oxygen consumption, and the obtained values were corrected for ROX.

## Acknowledgments

The authors thank Prof. Thomas Langer for providing the OMA1 KO cell line.

This work was supported by the Foundation for Polish Science First TEAM Programme co-financed by the European Union under the European Regional Development Fund (POIR.04.04.00-00-3F36/17); the National Science Centre Poland (NCN) grant (2019/34/E/NZ1/00321) in SONATA BIS Programme; the “Regenerative Mechanisms for Health - ReMedy” project (MAB/2017/2) carried out within the International Research Agendas Programme of the Foundation for Polish Science co-financed by the European Union under the European Regional Development Fund; the National Science Centre, Poland [2021/40/C/NZ3/00283]. Proteomic measurements were performed at the Proteomics Core Facility, IMol Polish Academy of Sciences. TEM analyses were performed at the Laboratory of EM of the Nencki Institute using infrastructure supported by EuBI Polish Node “Advanced Light Microscopy Node Poland”.

For the purpose of Open Access, the authors have applied a CC-BY public copyright license to this submission.

The authors declare no competing financial interests.

## Author contributions

P.Bragoszewski designed the study, performed experiments, and analyzed data. M.Krakowczyk performed live-cell experiments, and Western Blots analysis. A.Lenkiewicz performed experiments including confocal microscopy. T.Sitarz generated and characterized cell lines and contributed to the mass spectrometry experiment. D.Malińska processed confocal microscopy images. A.Wydrych carried out part of the fractionation experiments. A.Szczepankiewicz and H.Nieznanska prepared transmission electron microscopy images and statistics. B.Mussulini performed oxygen consumption rate experiments and analysis. R.Serwa and V.Linke acquired and analyzed mass spectrometry-based proteomics data. P.Bragoszewski and A.Chacinska provided funding. M.Krakowczyk assembled figures with the input of V.Linke, D.Malińska, B.Mussolini, and P.Bragoszewski. P.Bragoszewski and A.Krakowczyk wrote the manuscript with the input of all authors.

