## Supplementary Figures 1-3 for "OMA1 protease eliminates arrested protein import intermediates upon depolarization of the inner mitochondrial membrane"

**Supplementary figures and legends**

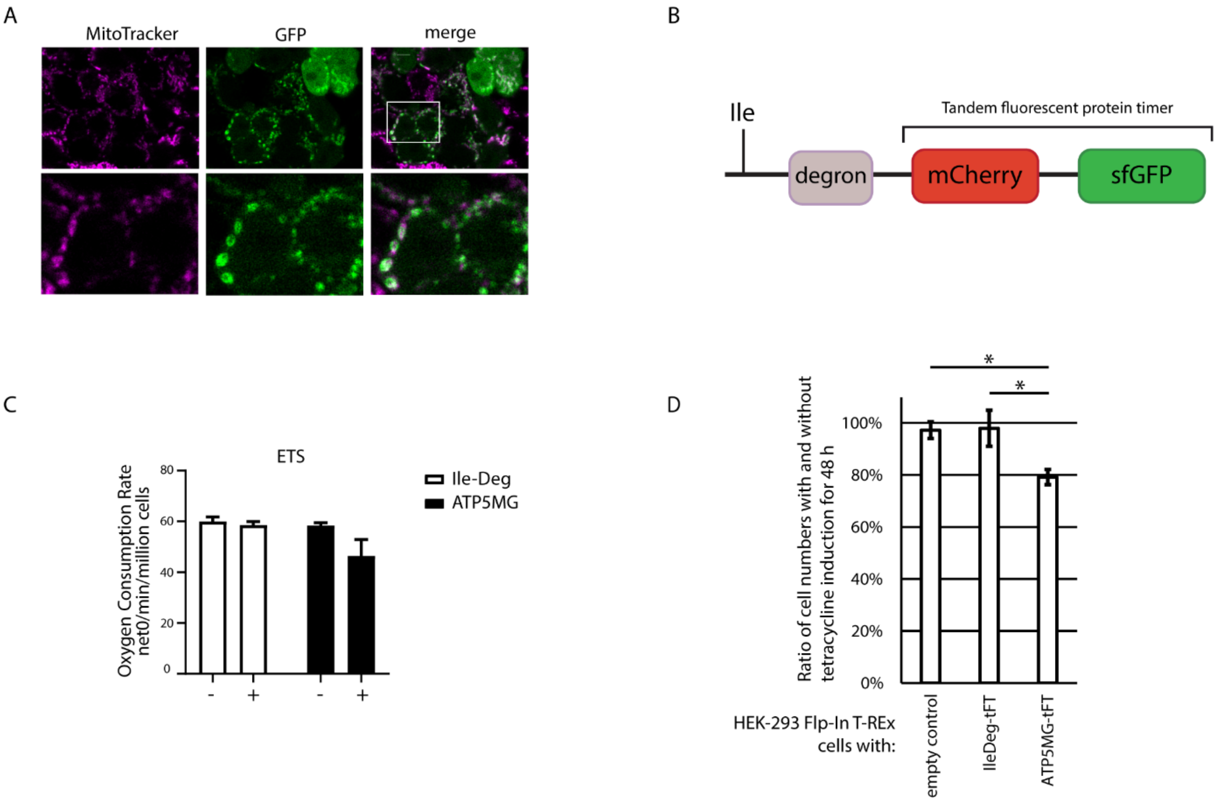


Figure S1 **Related to Figure 1.** (A) Confocal imaging of HEK-293 cells with ATP5MG-tFT fusion. Cells were live-stained with Mitotracker Deep Red FM. ATP5MG-tFT was detected by GFP fluorescence. Bottom panels show enlarged fragments marked in the top panel. (B) A schematic representation of Ile-Deg-tFT fusion protein (Khmelinskii et al., 2012). N-degron tFT fusion proteins are encoded as pro-N-degron with a N-terminal ubiquitin. Ubiquitin is removed from the translated protein exposing the new N-termini – isoleucine (Ile) amino acid residue was used in this work. Isoleucine was shown to confer a moderate turnover rate of the fusion (Khmelinskii et al., 2012). (C) Electron transport system (ETS) respiration based on Fig 1H. OCR was normalized to number of cells. Data are presented as the mean ± SEM ($n = 3$); *$p < 0.05$ (two-way ANOVA). (D) HEK-293 Flp-In T-REx cells with ATP5MG-tFT, IleDeg-tFT, or empty were cultured with or without addition of 1 µg/mL tetracycline for 48 h. Cell proliferation was measured by direct cell counting. Graph presents number of cells following induction as percentage of corresponding cells not treated with tetracycline. Mean ± SEM ($n = 4$). *$p < 0.05$ (Student's t-test).

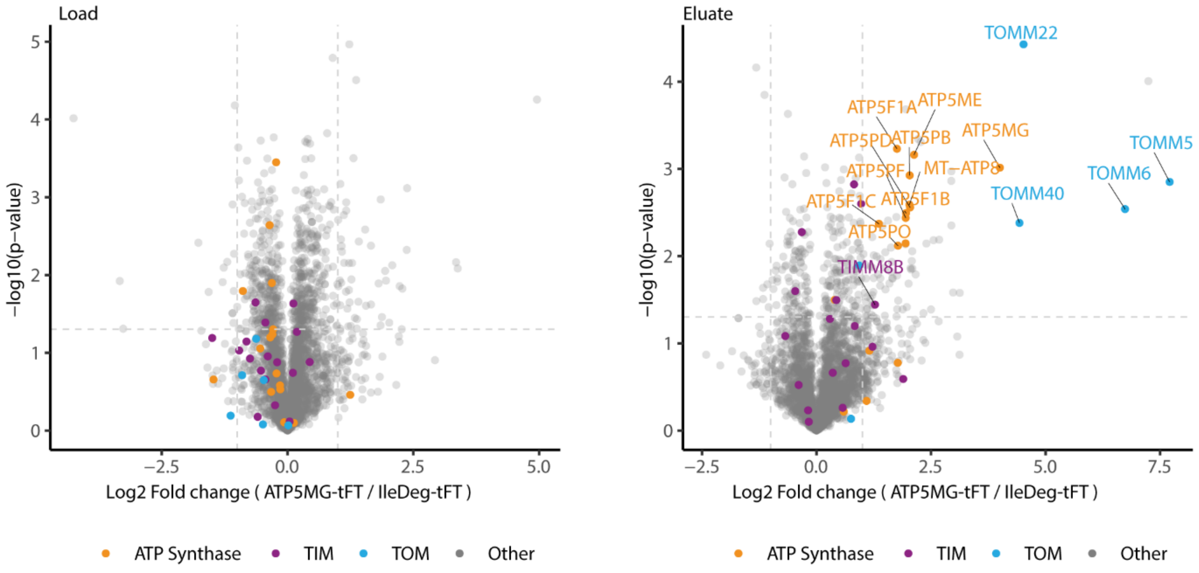


Figure S2 **Related to Figure 2.** Proteins from cellular lysates (Load) and purified with anti-GFP beads (Eluate) were analyzed by LC-MS/MS (n=4). Volcano plots x-axis represent $\log_2$ fold change of protein levels in Lysates and Eluates of HEK293 cells with ATP5MG-tFT compared to cells with IleDeg-tFT fusion proteins. Orange, magenta, and blue indicate proteins belonging to ATP synthase, TIM, and TOM complexes, respectively; significantly changed (Student's t-test $p<0.05$, $|\log_2$ fold change$| >1$) were labeled.

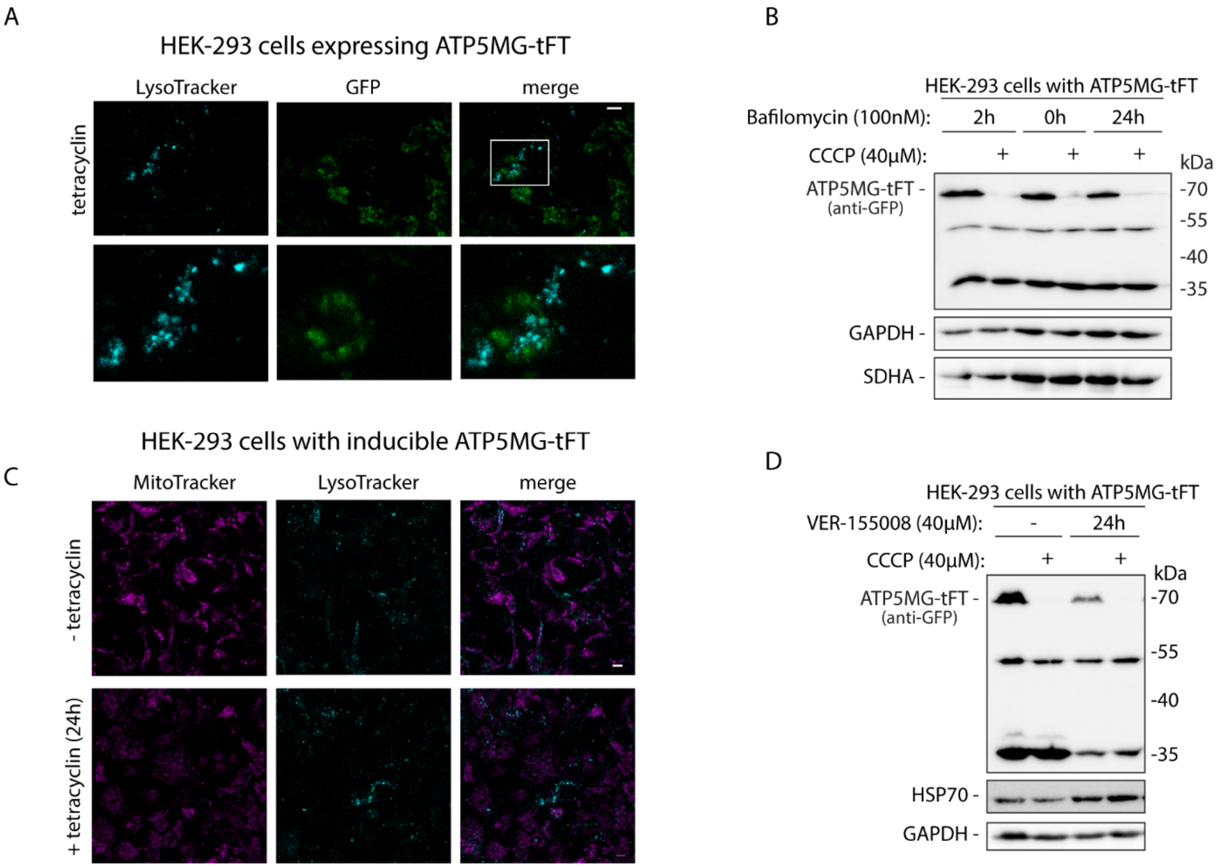


Figure S3 **Related to Figure 4.** (A) HEK-293 cells expressing ATP5MG-tFT were live-stained with LysoTracker and imaged by confocal microscopy. Scale bar 5 µm. Bottom panels show enlarged fragment marked in the top panel. (B) ATP5MG-tFT clogger levels in HEK-293 cells are not affected by autophagy inhibition using bafilomycin (100 nM). (C) Confocal imaging of HEK-293 cells with or without 24 h tetracycline induction of ATP5MG-tFT expression. Cells were live-stained with LysoTracker and Mitotracker Deep Red FM. Scale bar 5 µm. (D) HEK-293 cells with ATP5MG-tFT expression were treated with CCCP in the presence or absence of Hsp70 inhibitor, VER-15508 (40µM). Inhibition of HSP70 decreased the overall amount of ATP5MG-tFT clogger and its fragments, but did not affect CCCP induced processing.
